# Cognition Disconnected: The Influence of Domain-General Cognition (g) in Lesion-Deficit Mapping

**DOI:** 10.1101/2025.11.25.690275

**Authors:** Lara Sozer, Joseph Griffis, Kathleen Langbehn, Joel Bruss, Emily Olafson, Kenneth Manzel, Maurizio Corbetta, Daniel Tranel, Aaron D. Boes, Mark Bowren

## Abstract

Lesion-deficit mapping is a premier technique for dissociating neural correlates of cognitive functions. However, the contribution of domain-general cognition (g) to the performance of virtually all complex behavioral tasks may encumber such efforts. This possibility has been largely unexplored. Here, we examined the influence of g in lesion-deficit mapping using a combination of structural equation modeling, lesion-behavior mapping (LBM), and structural (sLNM) and functional (fLNM) lesion network mapping. We modeled latent variables for visuospatial ability, processing speed, language, and anterograde memory among 473 patients with focal brain damage (age range 20-93, 257 men, 216 women), both with and without g-associated variance embedded within each factor. Lesion-deficit maps for domain-specific cognitive abilities were statistically significantly less spatially inter-correlated after partitioning out g (*p* = 0.010 for LBM; *p* = 0.003 for sLNM; *p* = 0.003 for fLNM). Lesion-deficit maps also implicated the existence of distinct neuroanatomy for specific cognitive domains that was, in many cases, only apparent and clearly distinct from lesion-deficit maps of g after partitioning out the influence of g. Overall, our findings suggest that, if left unaccounted for, domain-general cognitive variance could tangibly affect conclusions about the extent to which different cognitive functions rely on distinct neuroanatomy.

---

A central goal of cognitive neuroscience is to delineate the architecture of cognition in a manner that reflects brain structure and function. The lesion method is a critical tool towards this end, offering not only a window into causal brain-behavior relationships, but also a probe with which to analyze the organization of mental faculties (Thiebaut de Schotten et al. 2020). Lesion-deficit mapping is often used to identify dissociable neural correlates of different cognitive processes. However, it is often the case that such analyses also yield substantial overlap for purportedly distinct abilities (Barbey et al. 2014; Biesbroek et al. 2014, 2021; Na et al. 2022; Moore et al. 2024). One potential explanation for this phenomenon is that higher-order cognitive systems play a role in task performance that is shared across domains. At any given level of analysis, more “domain-general” cognitive systems may interfere with our ability to identify unique neural correlates of more “domain-specific” abilities (Campbell and Tyler 2018; Morales et al. 2018; MacGregor et al. 2022; Bhaya-Grossman et al. 2025). Domain-general cognition (or general intelligence, g) lies at the top of the hierarchy, and could influence our ability to infer neural correlates of virtually any specific cognitive function (Spearman 1904). Thus, understanding the impact of g on lesion-deficit mapping of domain-specific cognitive functions might prove to be key to accurately inferring many causal brain-behavior associations.

There is converging evidence from studies of neurologically healthy individuals that supports this notion. Psychometrically, a latent g factor alone can account for nearly half of the inter-individual variation in performances across diverse cognitive tests (Deary et al. 2010; Dubois et al. 2018). Moreover, a majority of the reliable variance of many latent cognitive factors can be attributed to g (Watkins 2017). These behavioral findings are consistent with observations from the functional neuroimaging literature. Specifically, in addition to demonstrating spatially distinct patterns of brain activity for different functions, performance of diverse cognitive tasks has been associated with activity in a shared set of brain regions. These regions are thought to reflect the recruitment of domain-general cognitive resources (Duncan 2010; Niendam et al. 2012; Camilleri et al. 2018; Assem, Blank, et al. 2020; Assem, Glasser, et al. 2020; He et al. 2021), and have been associated with g or cognitive abilities closely linked to g (Colom et al. 2006; Jung and Haier 2007; Deary et al. 2010; Gläscher et al. 2010; Woolgar et al. 2010; Barbey et al. 2012; Assem, Blank, et al. 2020; Cipolotti et al. 2023). Altogether, this literature implies that attempts to map any specific cognitive domain to the brain may identify neuroanatomical correlates of both the domain of interest and of g without clearly disentangling the two. However, whether and how g exerts such an influence has remained largely unexplored. Without this knowledge, the veracity of psychometric boundaries among higher-order cognitive functions and their unique anatomical correlates will remain unclear.

The present study investigated the impact of g on lesion-deficit mapping of other cognitive abilities: visuospatial ability, learning/memory, processing speed, and language. Using behavioral data from a large cohort of patients with focal brain damage (n = 473), we performed structural equation modeling to derive factor scores for each domain-specific ability. We used a *correlated factors* approach to model each domain-specific cognitive ability with variance attributable to g embedded within its corresponding latent factor. We also used a *bifactor model* to model each of these abilities with g variance partitioned out of the domain-specific factors. We then used lesion-behavior mapping (LBM), and structural (sLNM) and functional lesion network mapping (fLNM) to link each cognitive factor to regional and network-level brain damage. By comparing corresponding maps from the correlated factors and bifactor models, we inferred whether and how g influenced the results. Figure 1 presents a schematic of the study design.

**Figure 1.**
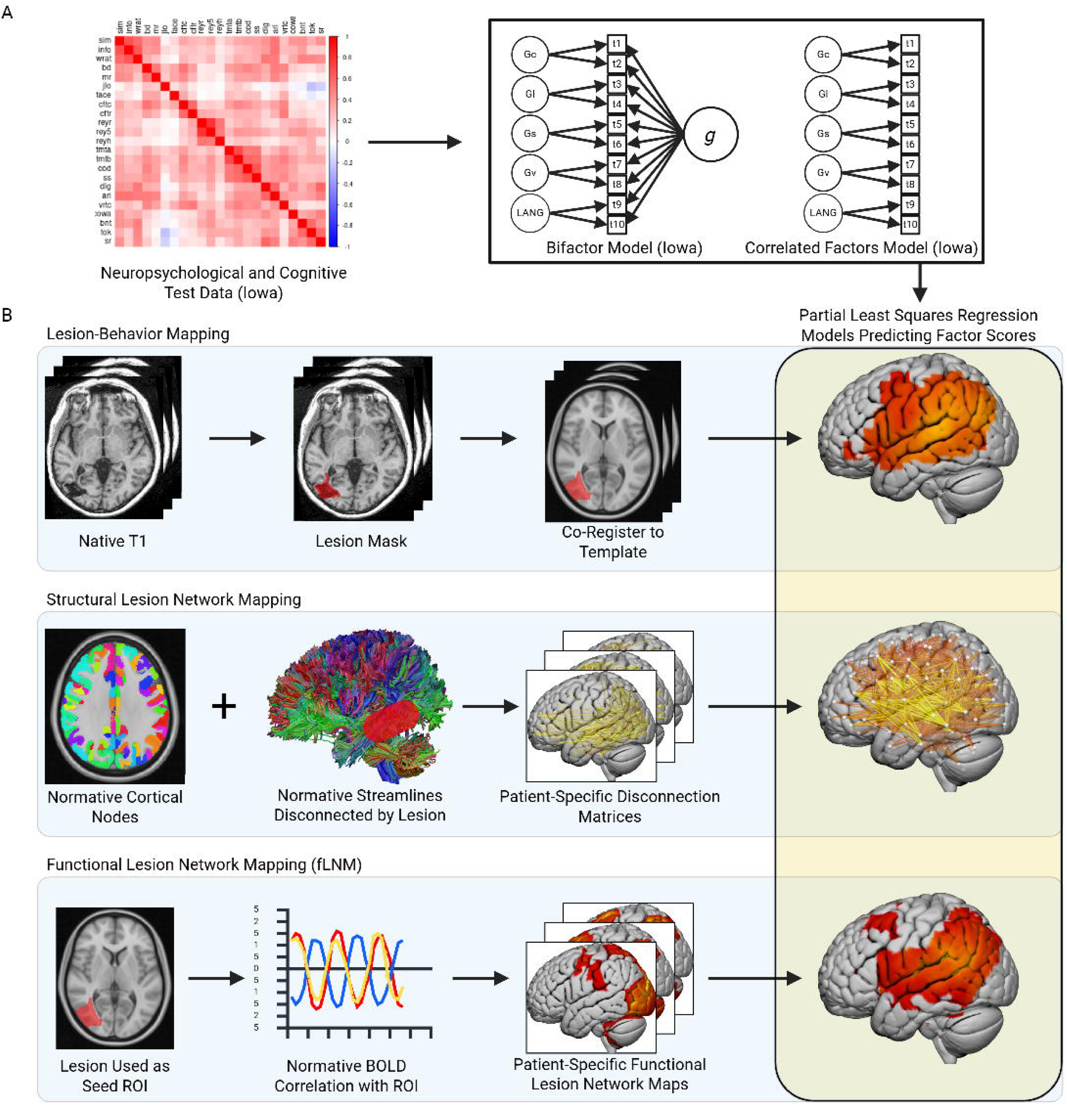
Study Design. (**A**) Neuropsychological and cognitive test data were used to model domain-general cognition (g) and domain-specific cognitive factors with structural equation modeling. We created a correlated factors model and a bifactor model to create latent factors for domain-specific abilities with and without g variance embedded in each factor, respectively. (**B**) Lesion-deficit mapping was used to link factors from the psychometric models to lesion location, white matter disconnections, and lesion-associated functional connectivity.

## Materials and Methods

### Participants and behavioral and neuroimaging data

We analyzed data from a total of 473 patients with focal brain damage (Tranel 2009) enrolled in the Iowa Neurological Patient Registry (Table 1). All participants gave written informed consent to participate in the studies described below, which were approved by the Institutional Review Boards of the University of Iowa.

**Table 1.**
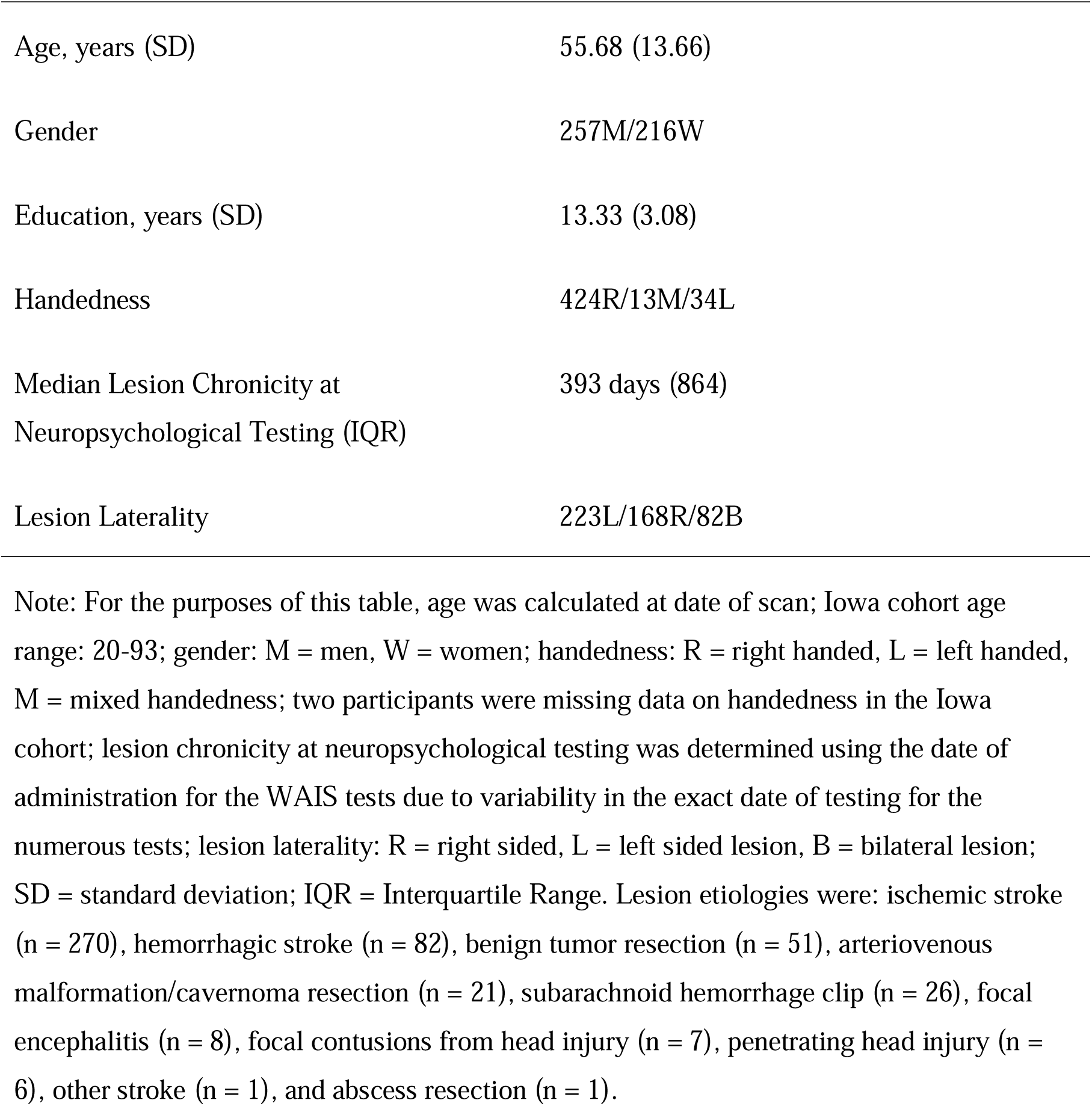
Demographic Information.

Neuropsychological and neuroimaging data were collected from participants three months or more after lesion onset. We focused on a subset of participants who completed at least 66% of the 23 neuropsychological and cognitive measures that are most commonly administered in the Iowa Registry protocol (Anderson and Tranel 2016). Supplementary Table 1 lists all tests. Notably, several of these measures are subtests from the Wechsler Adult Intelligence Scale (WAIS), which is the most widely used clinical test of intelligence and its facets. All neuropsychological and cognitive tests were corrected for demographic factors (e.g., age, education) based on normative data from associated test administration manuals or Benton Neuropsychology Clinic data as in prior work (Bowren et al. 2020, 2022). Participants had a lesion mask registered to the MNI ICBM152 T1 nonlinear 6^th^ generation asymmetric template. Unlike prior analyses of g based on the Iowa Registry, we used all commonly administered cognitive and neuropsychological tests, which resulted in the inclusion of 7 additional tests and 71 additional participants compared to prior studies. Many of the new participants were newly enrolled. Exclusion criteria were: major psychiatric illness, developmental intellectual disability, progressive neurological disease, lesion onset at less than 18 years of age, and developmental-onset epilepsy.

### Structural equation modeling

We created structural equation models using lavaan software (Rosseel 2012) in R version 4.4.1. Missing data were imputed using Full Information Maximum Likelihood. This was required for 20.8% of the total data, which is within an appropriate range for imputation based on prior empirical work (Junaid et al. 2025). We began by creating a bifactor model of cognitive abilities. In a bifactor model, all tests load onto a domain-general factor (g) and potentially also one or more domain-specific factors. Latent variables are also orthogonalized to one another (i.e., covariances forced to zero). We started with the bifactor model because this model fit best in prior analyses based on a smaller subset of the Iowa Neurological Patient Registry (Bowren et al. 2020). We then modified the baseline model to test for whether the inclusion of additional parameters improved model fit in terms of a Chi-Squared (Χ^2^) difference test (alpha level = 0.05), as well as the following fit indices: Robust Comparative Fit Index (CFI), Robust Tucker-Lewis Index (TLI), Robust Root Mean Square Error of Approximation (RMSEA), and Standardized Root Mean Square Residual (SRMR). If a more complex model provided a better fit to the data (i.e., the model with more estimated parameters), we used the more complex model going forward. Otherwise, we retained the simpler model. We tested for the added value of non-zero covariances between observed and latent variables, and hypothesized factor cross-loadings. The final, best-fitting model was used for subsequent lesion-deficit mapping analyses. Lastly, we reported the model-based reliability estimates (omega hierarchical) for each domain-specific factor because these are often of interest for psychometric purposes. To create a correlated factors counterpart to the bifactor model, we removed the orthogonality constraints on the latent variables and all g loadings (i.e., only domain-specific factors were modeled, and all were allowed to be inter-correlated). See the Supplementary Material for additional information on modeling procedures.

For presenting the modeling results in a figure, we used abbreviations for the domain-specific factors derived from the CHC model of cognitive abilities (Schneider and McGrew 2018a); otherwise, for clarity in the main text, we refer to each factor by its full name. The CHC model has been shown to fit neuropsychological test data well (Jewsbury et al. 2017; Schneider and McGrew 2018b).

### Lesion segmentation

Lesion boundaries were manually segmented in three dimensions using MRI as described previously (Bowren et al. 2020), or, in rare cases, a CT was used if MRI contraindications were present (n = 80 CT scans). A neurologist blinded to the behavioral data evaluated the anatomical accuracy of each lesion tracing, both before and after the transformation to the template brain. These evaluations were performed using a combination of linear and nonlinear registration techniques as described previously (Bowren et al. 2020, 2022). The final output for each patient was a binary, 3-dimensional volume indicating the presence or absence of the lesion. These were then down-sampled to 3 mm resolution to reduce downstream computational demands and because the meaningful resolution of lesion-associated cognitive deficits is likely not at the level of a single millimeter (Pustina et al. 2018).

### sLNM

We used the Lesion Quantification Toolkit (Griffis et al. 2021) to estimate the pairwise cortical regions disconnected by each patient’s lesion based on normative diffusion-weighted imaging-based tractography (Yeh 2022). This toolkit applies uses DSI Studio (https://dsi-studio.labsolver.org/) to identify normative streamlines from the HCP-1065 atlas that intersect each patient’s lesion volume. We also identified the normative group-average gray matter parcels interconnected by these streamlines, and calculated the proportion of those streamlines that intersected the lesion mask. Gray matter parcels were pulled from the Schaeffer atlas (Schaefer et al. 2018), Freesurfer DKT atlas, Cerebellum-SUIT atlas (Diedrichsen 2006), and the Harvard-Oxford atlas (https://fsl.fmrib.ox.ac.uk/fsl/fslwiki/Atlases). For each participant, the end result used in subsequent analyses was a 245 x 245 matrix. Within this matrix, each cell contained the proportion of streamlines connecting two parcels that were affected (i.e., estimated to be “disconnected”) by the lesion.

### fLNM

Brain networks affected by a lesion can also be estimated through normative resting-state fMRI connectivity (Boes et al. 2015; Fox 2018). We used the Brain Genomics Superstruct Project (GSP 1000) dataset (Cohen et al. 2020) to perform resting-state fMRI connectivity analyses with each patient’s lesion volume as a seed region-of-interest. As described previously, this involved correlating the average time course of blood oxygen level-dependent (BOLD) signal within the lesion volume with the BOLD signal observed in all other voxels of the brain (Bowren et al. 2022). We used principal component analysis to identify the first principal network component within each lesion volume (Pini et al. 2021). This network component was then used to derive a single voxel-wise map of the normative functional connectivity (represented as T values) associated with the lesion volume. This map was used in subsequent analyses.

### Statistical analyses

LBM, sLNM, and fLNM were performed using the Iowa Brain-Behavior Modeling (IBBM) Toolkit (Griffis et al. 2024). We used both mass-univariate and multivariate approaches as suggested by others (Ivanova et al. 2021). We used partial Pearson correlations controlling for log lesion volume for the mass-univariate models, and partial least squares regression (PLSR; also controlling for lesion volume through linear regression-based residuals) for the multivariate approach. PLSR depends on the model hyper-parameters (i.e. number of PLS components) and may not directly reflect the observed patterns of bivariate associations in the data. For this reason, the mass-univariate models are easier to interpret when the goal is to study voxel-wise brain-behavior associations using lesion data. Because our goal was to study how different cognitive modeling approaches affect inferences about brain-behavior relationships, we focused on the mass-univariate results. We did not map crystalized intelligence because it is thought to be largely robust to brain damage, and has not localized well in prior work (Bowren et al. 2020). Finally, spatial correlations between maps were performed in MATLAB using the corr function with the input being unthresholded *t*-statistics from the mass-univariate results (or the unthresholded PLSR model weights for the supplementary PLSR versions of the results). We used unthresholded maps for spatial correlations in order to ensure that all maps had the same number of voxel values and to avoid pitfalls of using thresholds (Sperber and Dadashi 2020). To statistically test for a difference in the spatial correlations among the maps from the correlated factors and bifactor models (i.e., to test whether bifactor-based maps were less spatially inter-correlated), we first converted the spatial correlations to Z scores using the Fisher r-to-Z transform, and then used permutation testing (with 10,000 permutations) to evaluate the statistical significance of a dependent samples *t*-test.

### Mass-Univariate Models

Mass-univariate analyses used Pearson correlations (or partial correlations when a confound, such as lesion volume, is specified in the analysis) between the input data and the factor scores. A single, brain-wide threshold for statistical significance was determined using permutation testing (1000 permutations) according to the previously published continuous family-wise error correction method (Mirman et al. 2018), which adjusts for multiple comparisons across the whole brain simultaneously (adjusted *p* threshold was set to 0.05).

The continuous family-wise error control method requires a “voxel count threshold”, and we used the most stringent voxel count threshold across all analyses for consistency (i.e., v = 1), which corresponds to the conventional whole-brain permutation correction approach (Mirman et al. 2018) and controls for the probability of at least one false positive voxel.

### PLSR models

We fit PLSR to the entire dataset at once after performing hyper-parameter optimization with cross-validation (5 folds, 5 repeats). We created separate PLSR models for the 3 predictor modalities: LBM (i.e., binary lesion volumes), sLNM (i.e., percent disconnection matrices), or fLNM (i.e., lesion-derived resting-state functional connectivity). Details on visualization of the PLSR model weights for supplementary figures are provided in the Supplementary Material. The variance explained by each PLSR model, calculated as the correlation between the predicted scores from the model fit to the full dataset and the observed scores, are presented in Supplementary Material.

### Data availability

The data that support the findings of this study are available on request from the corresponding author. The data are not publicly available due to privacy and ethical restrictions.

## Results

Demographic and lesion etiology data are presented in Table 1. The anatomical distribution of the lesions is depicted in Supplementary Figure 1. There was good lesion coverage of the cerebrum.

### Structural equation modeling

The final behavioral models are depicted in Figure 2. See the Supplementary Material for additional psychometric modeling results. The bifactor model converged normally and all factor loadings were statistically significant (Robust CFI = 0.948, Robust TLI = 0.933, Robust RMSEA = 0.055, SRMR = 0.054). Omega- hierarchical coefficients were: g (0.594), crystalized intelligence (0.335), verbal learning/memory (0.567), processing speed (0.450), visuospatial ability (0.587), and language (0.250). The same coefficients for the correlated factors version of this model (Robust CFI = 0.912, Robust TLI = 0.894, Robust RMSEA = 0.070, SRMR = 0.063) were: crystalized intelligence (0.618), verbal learning/memory (0.610), processing speed (0.712), visuospatial ability (0.745), and language (0.524).

**Figure 2.**
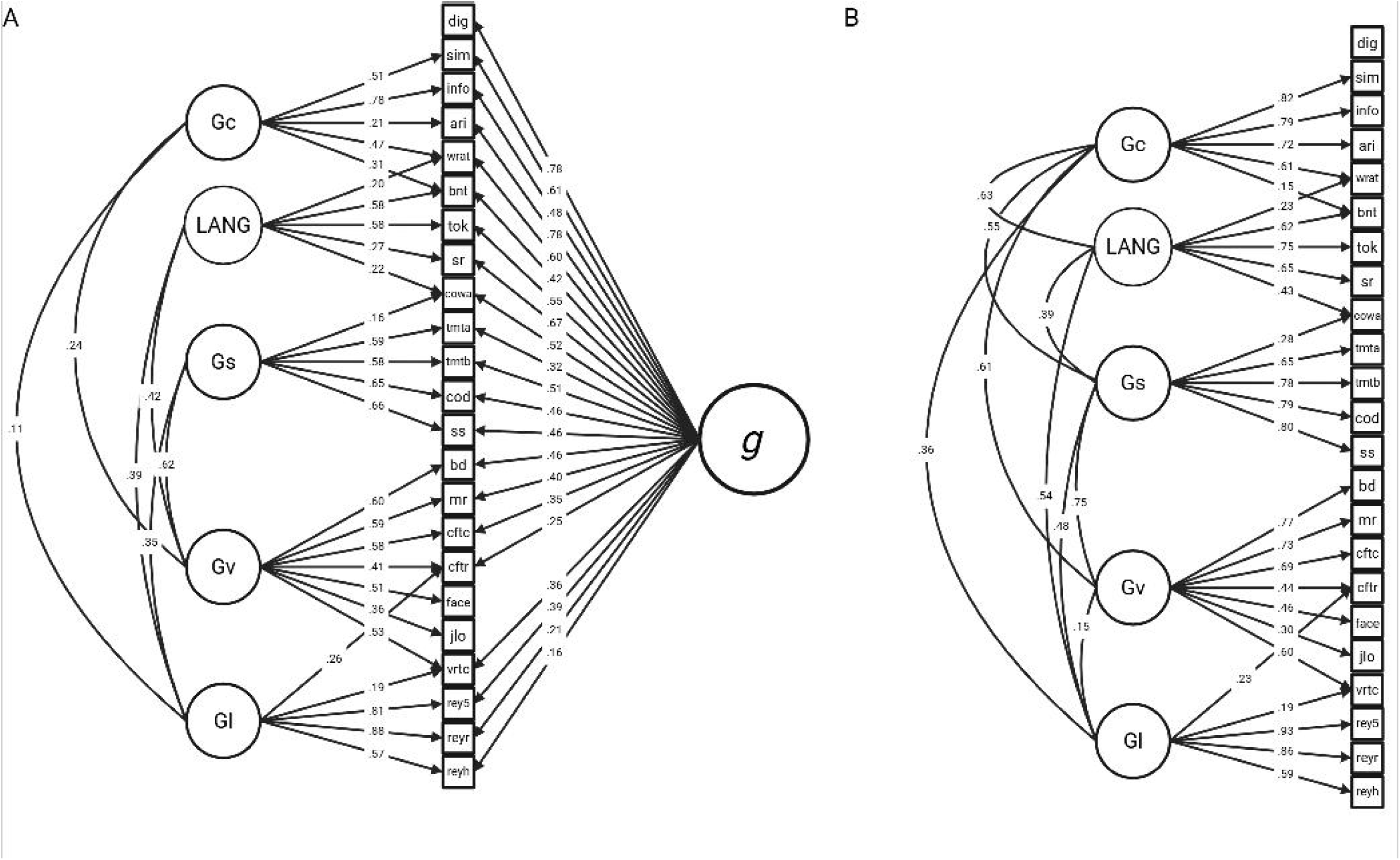
Structural Equation Modeling. (**A**) The final bifactor and (**B**) correlated factors structural equation models are depicted. LANG = language; Gs = processing speed; Gv = visuospatial ability; Gl = learning / memory; Glv = visuospatial learning / memory; Gla = auditory-verbal learning / memory; dig = WAIS Digit Span; sim = WAIS Similarities; info = WAIS Information; ari = WAIS Arithmetic; WRAT = WRAT Word Reading; bnt = BDAE Boston Naming Test; tok = MAE Token Test; sr = MAE Sentence Repetition; cowa = MAE Controlled Oral Word Association; tmta = Trail-Making Test Part A; tmtb = Trail-Making Test Part B; cod = WAIS Digit-Symbol Coding; ss = WAIS Symbol Search; bd = WAIS Block Design; mr = WAIS Matrix Reasoning; cftc = Complex Figure Test Copy Trial; cftr = Complex Figure Test Delayed Recall Trial; face = Benton Facial Recognition Test; jlo = Benton Judgment of Line Orientation Test; vrtc = Benton Visual Retention Test Total Correct; rey5 = RAVLT Trial 5; reyr = RAVLT Delayed Recall; reyh = RAVLT Delayed Recognition Hits; We omitted correlated residuals between observed variables (cftc and cftr, tmta and tmtb) for visual clarity.

### LBM, sLNM, and fLNM of g

LBM linked g to the left temporo-parietal-occiptal junction, left auditory association regions, and left hemisphere white matter (Figure 3). sLNM linked g to widespread disconnections in the left hemisphere, most strongly implicating disconnections between the left lateral prefrontal and lateral temporal cortices, between the left lateral temporal and parietal cortices, and the bilateral temporal and parietal cortices. Regions linked to g via fLNM included the left temporo-parietal occipital junction, left posterior superior temporal gyrus, left frontal operculum, and left posterior middle frontal gyrus. Controlling for lesion chronicity produced virtually identical results (Supplementary Figure 2).

**Figure 3.**
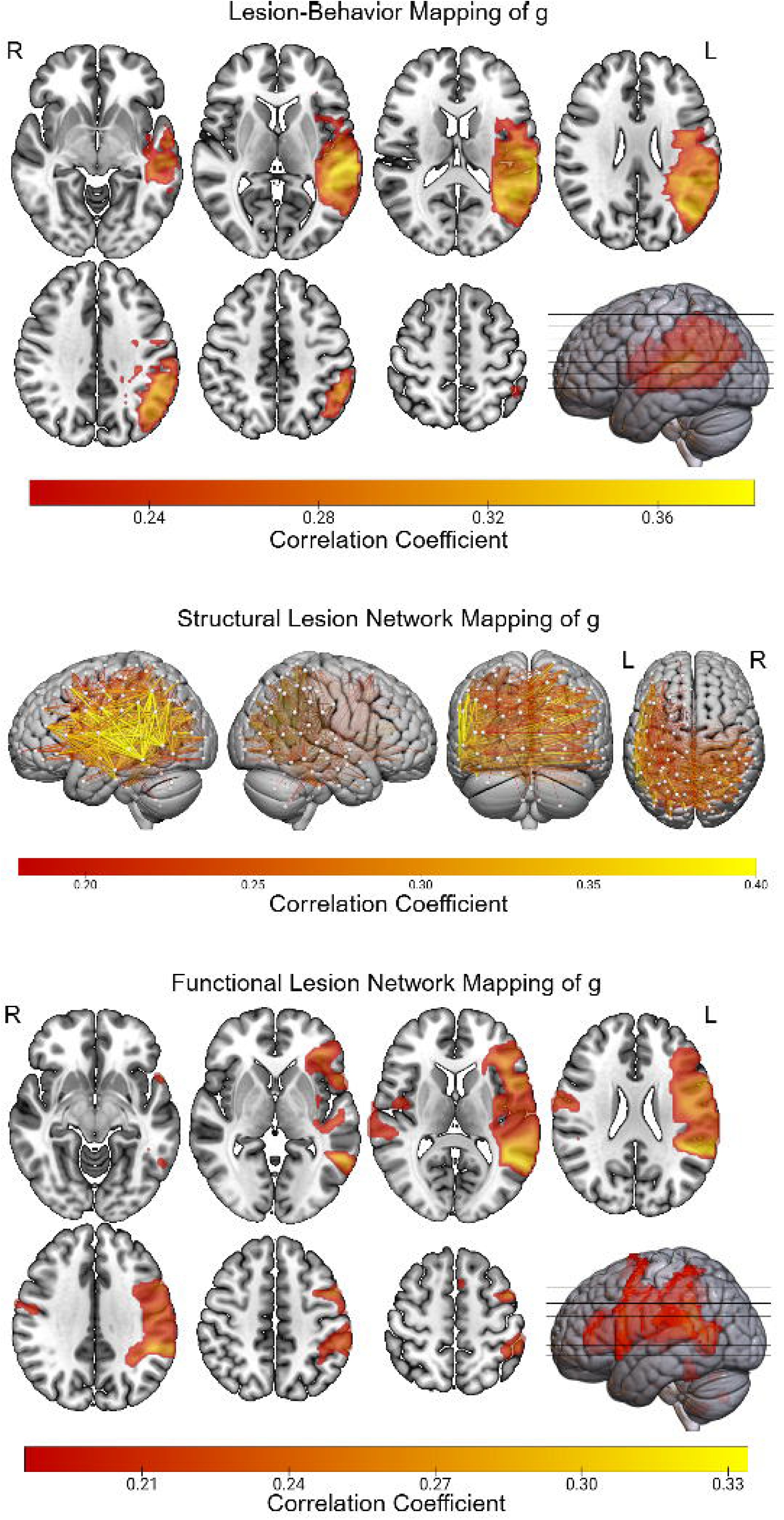
LBM, sLNM, and fLNM of. **g.** The LBM, sLNM, and fLNM of g are depicted. Voxel values (for LBM and fLNM) and edge values (for sLNM) indicate the Pearson correlation with g from the mass-univariate models after correction for multiple comparisons using brain-wide family-wise error.

### Spatial Correlations Among Lesion-Deficit Maps of Domain-Specific Cognitive Abilities are Lower After Controlling for g

We first evaluated whether spatial correlations among maps of domain-specific abilities were lower after controlling for g (i.e., in the bifactor model) compared to corresponding maps that did not account for g (i.e., in the correlated factors model; Figure 4). Across LBM (permutation *p* = 0.010), sLNM (permutation *p* = 0.003), and fLNM (permutation *p* = 0.003), permutation testing demonstrated that the spatial correlation values (calculated from unthresholded maps and then converted to Z scores as noted in the methods) from maps of domain-specific cognitive abilities were significantly lower for maps based on the bifactor model compared to those based on the correlated factors model.

**Figure 4.**
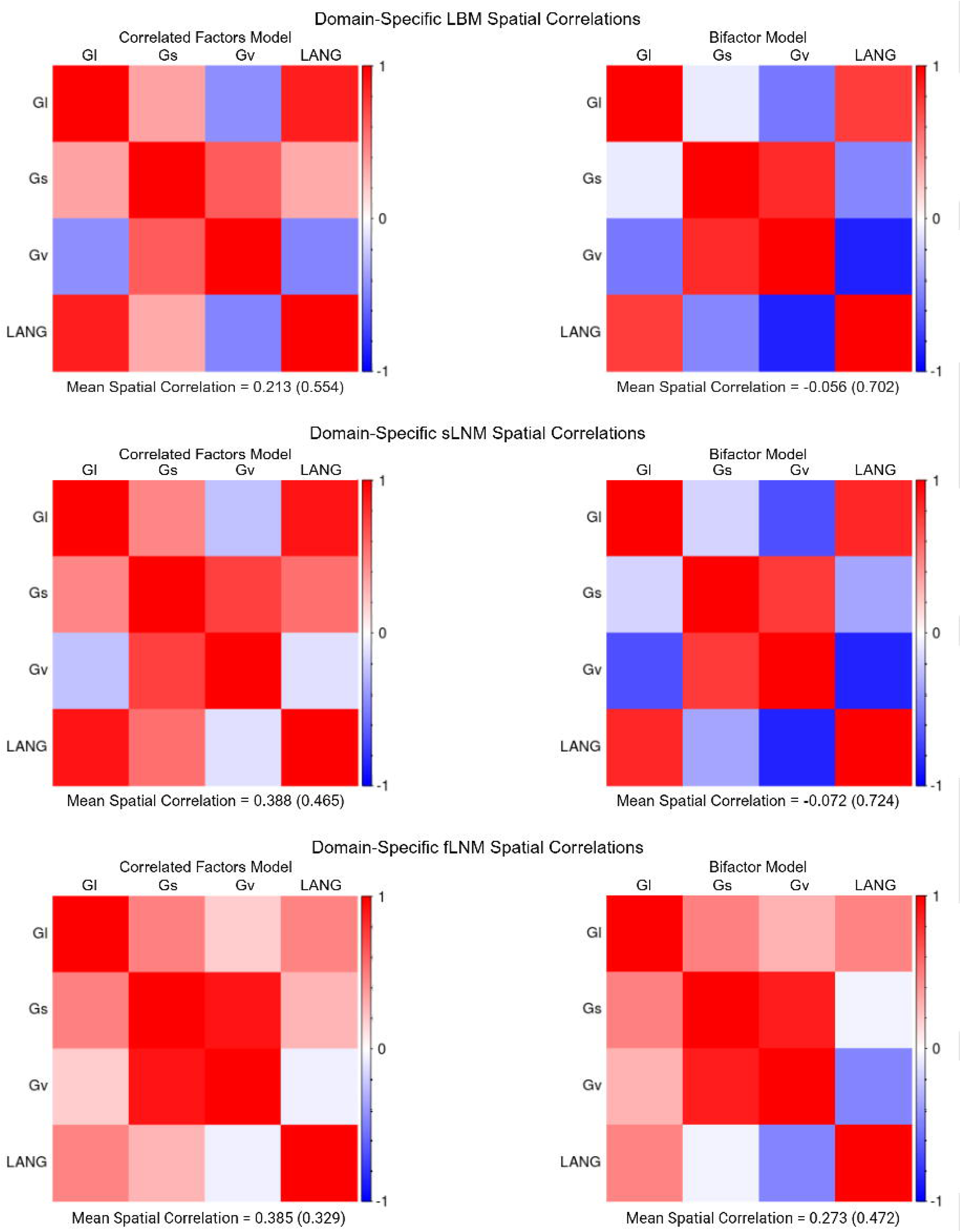
Spatial Correlations Among Domain-Specific Lesion-Deficit Maps with and without Controlling for. **g.** We correlated the spatial distribution of the unthresholded *t*-statistics from the mass-univariate models for the domain-specific cognitive factors (i.e., all non-g factors). We separately depict the spatial correlation matrices for LBM, sLNM, and fLNM based on the correlated factors and bifactor models. Across the 3 lesion-deficit mapping modalities (i.e., LBM, sLNM, and fLNM), the mean spatial correlation was higher for the maps from the correlated factors model compared to those derived from the bifactor model. Standard deviations are provided in parenthesis. Statistical comparison of the distribution of spatial correlations was performed after converting spatial correlations to Z scores using the Fisher’s r-to-Z transform. Permutation testing (with 10,000 permutations) of a dependent samples *t*-test on the Z scores indicated that the spatial correlations were statistically significantly lower for the bifactor maps compared to the correlated factors maps (*p* = 0.010), sLNM (*p* = 0.003) and fLNM (*p* = 0.003).

For each cognitive ability factor and each lesion-deficit mapping modality, we also calculated the spatial correlation between the unthresholded *t*-statistics from the bifactor model-based map and its correlated factors counterpart (Supplementary Table 3). The most dissimilar pairs of maps were the LBMs of processing speed (*r* = 0.588) and the sLNMs of processing speed (*r* = 0.692). This suggests that maps of processing speed were most strongly altered by g variance, which is notable given that g has long been linked to processing speed.(Schubert et al. 2017; Mashburn et al. 2024)

### Comparing Lesion-Deficit Mapping Results with Versus without Controlling for g

Anatomy associated with the LBMs, sLNMs, and fLNMs are presented in Supplementary Tables 4-6 (Glasser et al. 2016; Huang et al. 2022; Yeh 2022). Below, we focus on describing notable differences between the correlated factors- and bifactor-based mass-univariate results for each cognitive domain. Multivariate (i.e., PLSR-based) results of these analyses are depicted in Supplementary Figures 3-7 and in Supplementary Table 7.

### Visuospatial Ability

Mass univariate results for the visuospatial ability factors from the bifactor and correlated factors models are depicted in Figure 5. LBM of the bifactor version of visuospatial ability, but not its correlated factors counterpart, implicated right insula/sub-insula and associated white matter, and deep right parietal white matter. The sLNM of the bifactor version of the visuospatial ability factor more strongly implicated right hemisphere fronto-posterior disconnections, and less prominently featured inter-hemispheric disconnections (especially those involving the left lateral temporal and left inferior parietal regions) compared to the correlated factors version of these results. For fLNM, the bifactor-based results implicated many more regions than the correlated factors-based results. For example, the thalamus and left cerebellum appeared in the bifactor mapping but not in the correlated factors mapping. Also, the right occipito-parietal region was more robustly implicated by results for the bifactor model.

**Figure 5.**
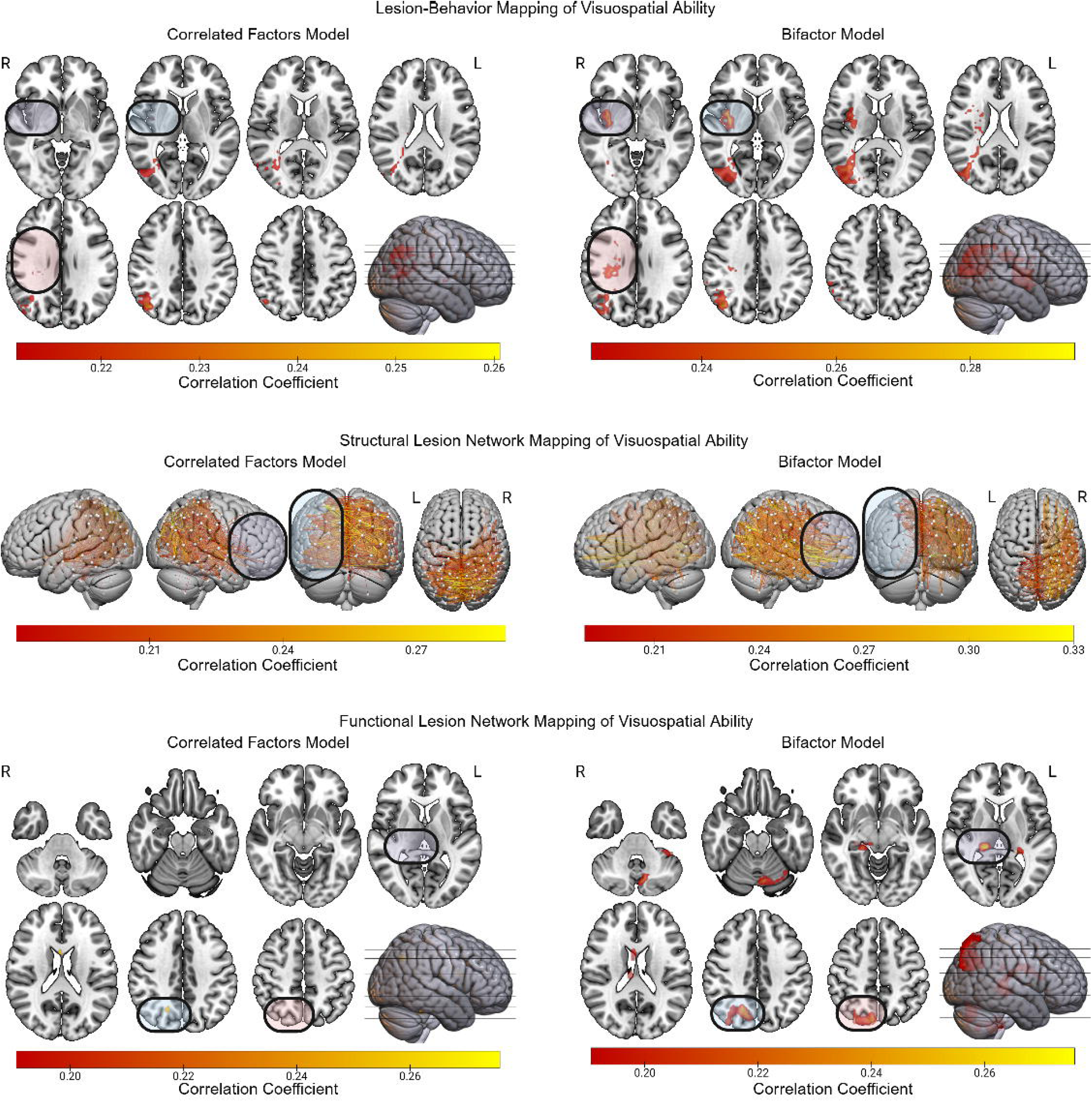
LBM, sLNM, and fLNM of Visuospatial Ability with and without Controlling for. **g.** The LBM, sLNM, and fLNM mass-univariate results are depicted for visuospatial ability factor scores derived from both the correlated factors model and the bifactor model. Differences between the results for the bifactor and correlated factors models are highlighted.

To illustrate the practical significance of these differences, we present case examples in which low visuospatial ability was observed following lesions involving regions linked to domain-specific visuospatial ability versus g (Figure 6). This shows how different lesion locations (and even in different hemispheres) can result in low visuospatial ability if g is not accounted for in the model. Both patients had low visuospatial ability factor scores from the correlated factors model, but for different reasons. The first patient had a right hemisphere lesion that affected anatomy linked to visuospatial ability by the bifactor results (cf. Figure 5 LBM from bifactor model). The second patient had a left hemisphere lesion that affected anatomy linked to g (cf. Figure 3 LBM). The bifactor model’s visuospatial factor was low *only* for the patient who had a lesion involving the right hemisphere anatomy linked to visuospatial ability.

**Figure 6.**
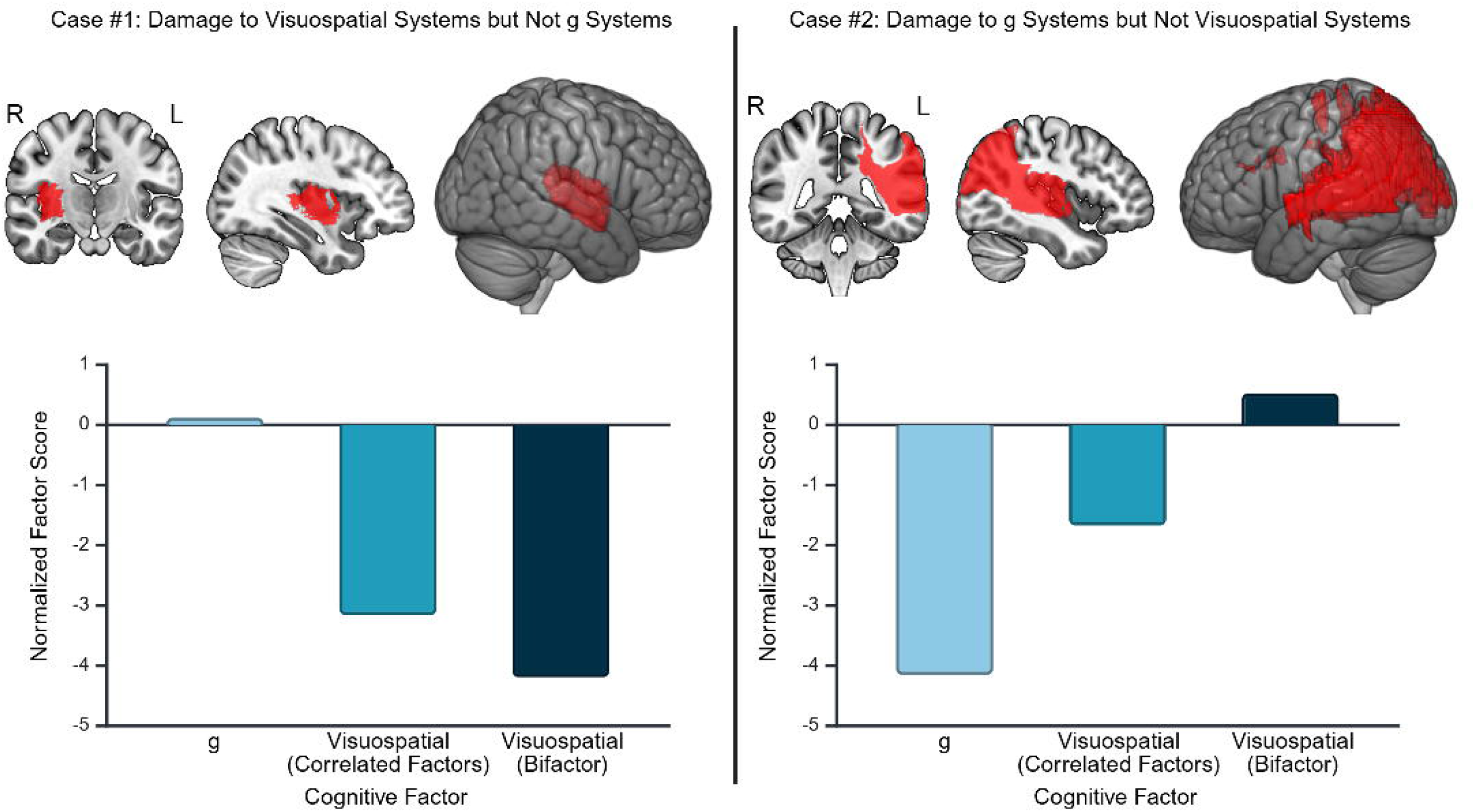
Case Examples Demonstrating Different Routes to Low Cognitive Factor Scores in the Correlated Factors Model. We depict case examples to illustrate how cognitive modeling can reveal different routes to impairment in visuospatial ability. In the first case example (left panel), we present data from a patient with a lesion involving a anatomy implicated by LBM of the bifactor model version of visuospatial ability (which contains variance due to only visuospatial ability), but not by LBM of its correlated factors counterpart (which contains variance attributable to both g and visuospatial ability). In the second case example (right panel), we present a data from a patient with lesion affecting anatomy linked to g but not to the bifactor version of visuospatial ability. The first patient exhibited low visuospatial ability in both the correlated factors and bifactor models, while also having average g. The second patient exhibited low visuospatial ability in the correlated factors model, but upon inspection of the bifactor model scores, it is apparent that this is due to low g despite intact visuospatial ability. Thus, lesion-deficit mapping based only on the correlated factors version of visuospatial ability would yield some results that could be attributable to an impairment in g rather than an impairment in visuospatial ability.

### Processing Speed

LBM linked the correlated factors version of processing speed to left hemisphere white matter deep to the temporo-parieto-occipital junction (Figure 7). Only a few scattered voxels were present in this region in the LBM of the bifactor version of processing speed. Nearly all findings for the LBM of the bifactor version of processing speed were located in the right hemisphere, including the right temporo-occipital region (region MST according to the HCP MMP1 extended atlas) and underlying white matter. Compared to the sLNM findings for the correlated factors version of processing speed, the bifactor-based results implicated far fewer inter-hemispheric disconnections involving left lateral temporal and inferior parietal cortices. The bifactor sLNM results largely implicated disconnections involving the occipital lobes.

**Figure 7.**
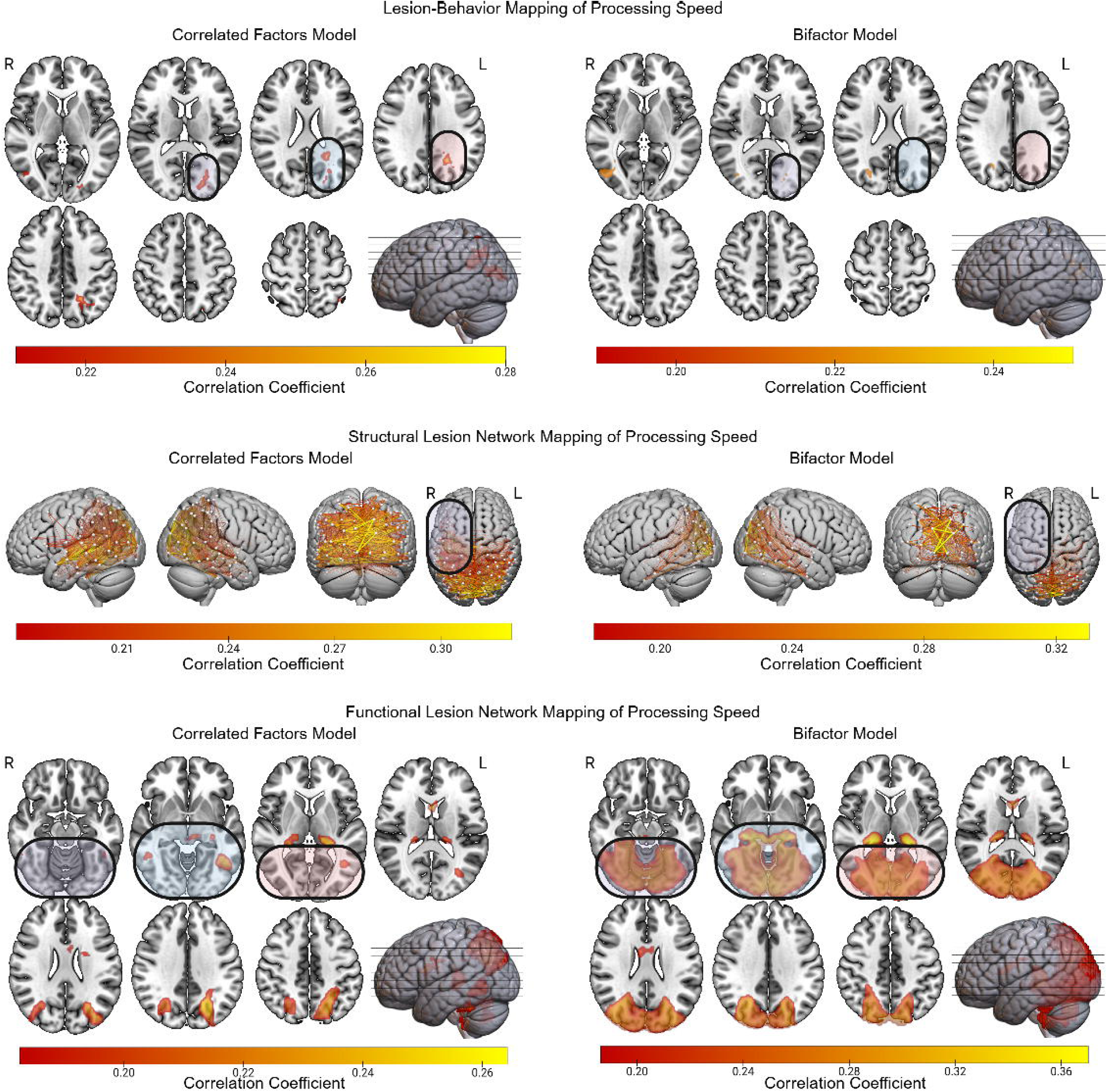
LBM, sLNM, and fLNM of Processing Speed with and without Controlling for. **g.** The LBM, sLNM, and fLNM mass-univariate results are depicted for Gs scores derived from both the correlated factors model and the bifactor model. Differences between the results for the bifactor and correlated factors models are highlighted.

The fLNM results based on the correlated factors model associated processing speed with a bilateral network involving posterior temporal, occipito-parietal, and thalamic regions. By contrast, the bifactor fLNM results implicated inferior occipito-temporal and occipito-parietal regions which most closely resemble the canonical visual network.

### Verbal Learning/Memory

The bifactor-based LBM of verbal learning/memory implicated only the left mesial temporal/inferior occipito-temporal region (Figure 8). By contrast, the correlated factors-based LBM results extended beyond this mesial temporal region to include the left lateral temporal cortex and underlying white matter, and the left parietal opercular region. Similarly, the sLNM results for the correlated factors version of verbal learning/memory implicated disconnections involving more superior regions (e.g., left lateral prefrontal cortex) that were not implicated in its bifactor counterpart. The fLNM based on the bifactor model implicated left mesial temporal and occipital regions. These regions were not linked to verbal learning/memory in the correlated factors version of the fLNM results.

**Figure 8.**
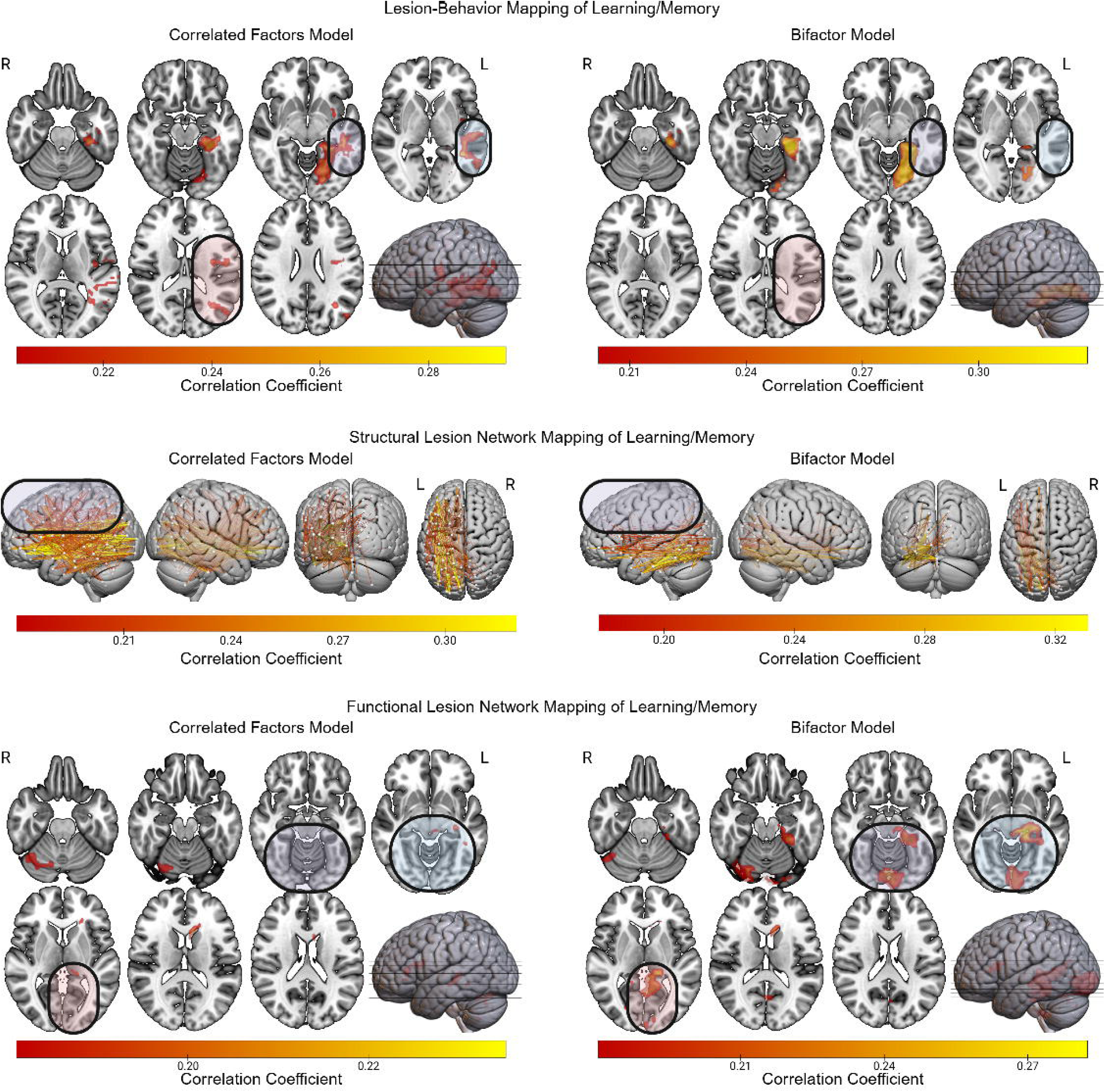
LBM, sLNM, and fLNM of Verbal Learning/Memory with and without Controlling for. **g.** The LBM, sLNM, and fLNM mass-univariate results are depicted for Gl scores derived from both the correlated factors model and the bifactor model. Differences between the results for the bifactor and correlated factors models are highlighted.

### Language

The LBM results for language based on the correlated factors model included more posterior regions than their bifactor counterpart (Figure 9). The bifactor-based LBM results also included a region in the medial parietal lobe that was not observed in the correlated factors version of the results. The sLNM based on the correlated factors version of language implicated a large number of disconnections involving the right lateral occipital, parietal, and temporal regions. By contrast, the sLNM based on the bifactor model did not include any of these disconnections. For fLNM, the correlated factors model version of language was linked to left posterior frontal cortex and left inferior sensorimotor cortex. Neither of these regions was implicated in the bifactor version of the fLNM of language, which was instead limited to the left inferior frontal and posterior temporal regions.

**Figure 9.**
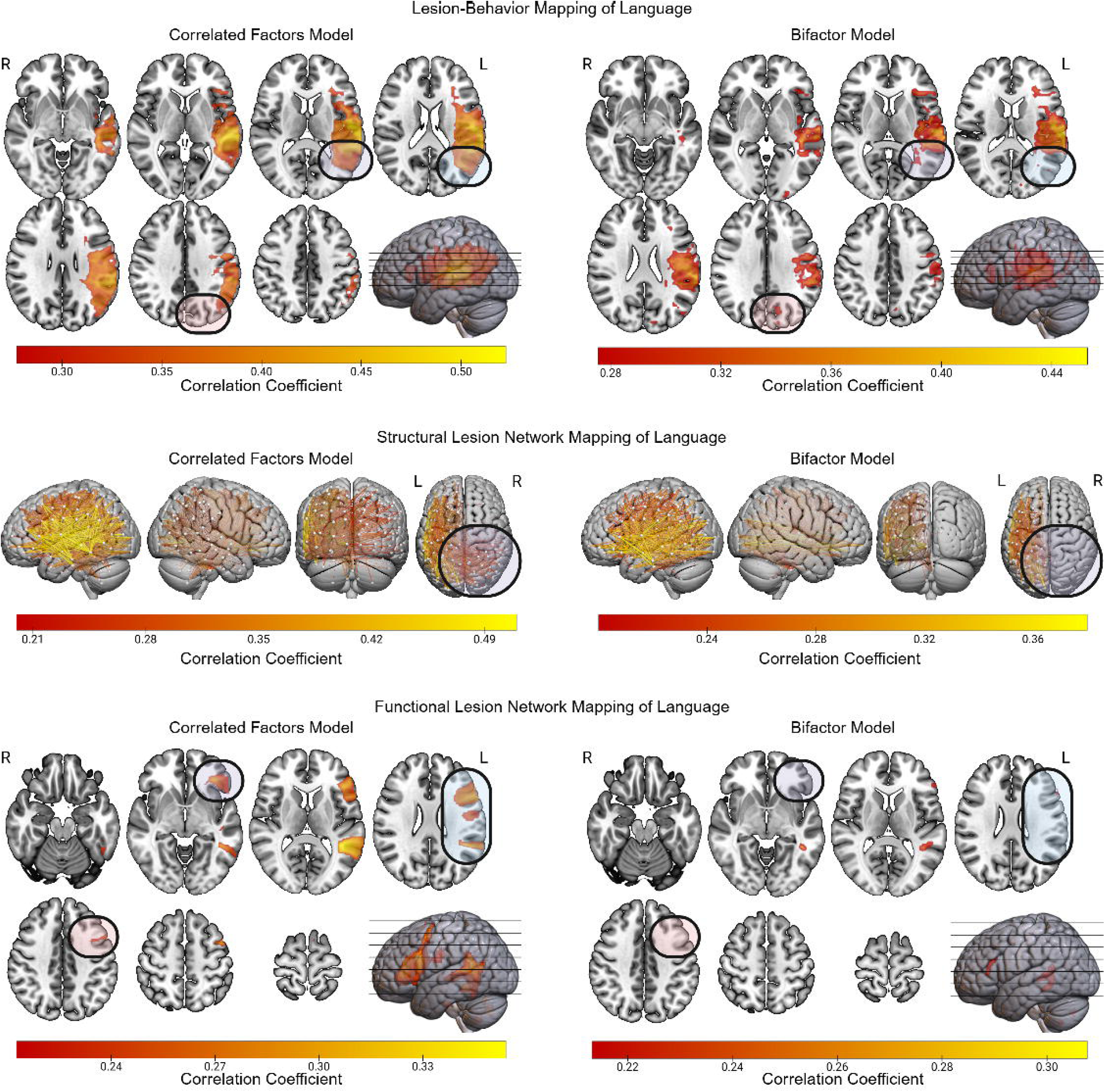
LBM, sLNM, and fLNM of Language with and without Controlling for. **g.** The LBM, sLNM, and fLNM mass-univariate results are depicted for LANG scores derived from both the correlated factors model and the bifactor model. Differences between the results for the bifactor and correlated factors models are highlighted.

### LBM and LNM of individual tests

Lastly, we performed LBM, sLNM, and fLNM on individual test scores to complement analyses based on cognitive factor scores (Supplementary Figures 7-10; see Supplementary Material for additional information on these analyses). Many of these results mirrored those based on factor scores. For example, LBM of Trial 5 from the Rey Auditory-Verbal Learning Test (a test of verbal learning/memory) identified left lateral posterior temporal regions that were also observed in the LBM of the verbal learning/memory factor from the correlated factors model. After controlling for g, these results were largely restricted to the inferior occipito-temporal/mesial temporal region, as was the case for the bifactor version of the verbal learning/memory factor.

## Discussion

With well over a century of foundational contributions to neuroscience, the lesion method has proven to be integral to efforts to dissociate human cognitive functions and their neural correlates. However, lesion-deficit mapping may be hindered by a problem of cognitive process specificity, which has remained largely unexplored. Functional imaging studies give reason to suspect that the entanglement of generalized and specialized systems for cognition affects conclusions about whether neural correlates of different cognitive abilities can be dissociated (Barbey et al. 2014; Biesbroek et al. 2014, 2021; Na et al. 2022; Moore et al. 2024). The present study demonstrates that this entanglement can impact lesion-deficit mapping, as we found that accounting for g altered lesion-deficit maps across cognitive domains and across lesion-behavior and lesion network mapping. In sum, this study highlights the importance of accounting for g in efforts to dissociate neural correlates of specific cognitive functions. Below, we discuss the implications of our findings, first with respect to g and how it broadly influenced the domain-specific lesion-deficit mapping results. We then discuss what can be gleaned from how g influenced the results for each specific cognitive domain. Lastly, we comment on important limitations of the present study.

Our results converge on the notion that fronto-temporo-parietal brain networks are key neural correlates of g. This is consistent with both functional imaging studies (Jung and Haier 2007; Camilleri et al. 2018) and lesion-deficit mappings of g (Gläscher et al. 2010; Barbey et al. 2012; Bowren et al. 2020) The present study extends on prior work by contributing the first explicit lesion network mapping analyses of g using both structural and functional brain network data. Consistent with prior lesion work by multiple groups (Barbey et al. 2012; Bowren et al. 2020), we found that g was principally linked to left hemisphere lesions (Deary et al. 2010; Bowren et al. 2020). This finding begs the question: is g truly a left lateralized cognitive ability, or is this finding driven by language demands embedded within virtually all cognitive and neuropsychological tests? Of note, the tests used to model g in this study included a roughly equal number of verbally mediated and visually mediated tests (Supplementary Table 1), which suggests that this finding was not merely driven by over-representation of verbally mediated tests. Moreover, our lesion-deficit mapping results of language from the bifactor model suggest that neural correlates of language are at least partially dissociable from g. However, it is important to note that our study was not designed to identify explanations for the left lateralization of g. Thus, further investigation into this topic is warranted. Regardless of why g has been linked to the left hemisphere, our results suggest that domain-general cognitive variance affects the anatomy implicated by lesion-deficit mapping of domain-specific cognitive abilities.

Across the non-g domains, there was a tendency for anatomy linked to g to be present in the correlated factors versions of LBM and LNM results, and yet absent in their bifactor analogues. For example, we observed this phenomenon with posterior inter-hemispheric disconnections in sLNM, and focal temporo-parietal white matter damage in LBM. These two findings were either less prominent or absent in lesion-deficit mappings of domain-specific functions after controlling for g via the bifactor model. These findings suggest that, at least in some circumstances, g-associated brain regions and networks can account for anatomical overlap in lesion-deficit mapping results for domain-specific cognitive functions. This is supported by our empirical comparison of the spatial correlations among the lesion-deficit maps based on the bifactor model versus those based on the correlated factors model. Specifically, we found that LBM, sLNM, and fLNM results based on the bifactor model were less similar to one another than those based on the correlated factors model. The present study thus suggests that accounting for domain-general systems could be important for isolating the anatomy unique to domain-specific cognitive functions.

Regarding the domain-specific lesion-deficit mappings, LBM of visuospatial ability after controlling for g implicated the right insula/sub-insula. This region contains white matter projections within the inferior fronto-occipital fasciculus. Thus, the LBM finding in this region may reflect the importance of right fronto-posterior disconnections for deficits in visuospatial ability. This conclusion is consistent with the sLNM findings for visuospatial ability, which more prominently implicate such disconnections only after controlling for g. These results implicate the interaction between brain regions for early visual representations and higher-order cognitive processing as central to visuospatial cognition (Mishkin et al. 1983; Heilman et al. 1986; Ungerleider et al. 1998; Wojciulik and Kanwisher 1999).

For processing speed, LBM based on the correlated factors model implicated a left hemisphere white matter region that we have previously linked to g (Bowren et al. 2020). After controlling for g, the LBM findings involved the right temporo-parieto-occipital region and underlying white matter, suggesting that the left-lateralization observed in the correlated factors-based result was due to the influence of g. The sLNM and fLNM findings based on the bifactor model converged on the importance of the bilateral occipitotemporal cortices/canonical visual network. These results are consistent with the fact that most of the processing speed tests involved visuomotor demands. They also suggest that amodal aspects of processing speed are tightly linked to g (Mashburn et al. 2024).

As for verbal learning/memory, controlling for g tended to focus lesion-deficit mapping results onto the left occipito-temporal/mesial temporal region. This result replicates one of the most robust findings in the field of cognitive neuroscience – the association of the mesial temporal lobe with anterograde memory (Scoville and Milner 1957; Zola-Morgan et al. 1986; Bartsch et al. 2011). By contrast, lateral prefrontal and posterior temporal regions, both of which were linked to g, were only implicated in verbal learning/memory when g was left unaccounted for in the model. These results echo neuropsychological studies suggesting that poor performances on tests of anterograde memory can be attributable to anterograde memory deficits proper, and/or executive dysfunction (Kopp and Thöne-Otto 2003; Zahodne et al. 2011). Our findings support this notion and suggest that the executive demands of memory tests may be linked to g.

Finally, with respect to language, controlling for g resulted in maps that were more anatomically restricted across LBM, sLNM, and fLNM. For example, in the sLNM results, inter-hemispheric disconnections were not implicated after controlling for g. Prior work has shown that the Multiple-Demand Network and a network linked to language processing are anatomically adjacent, yet distinct (Fedorenko et al. 2012; Campbell and Tyler 2018; Diachek et al. 2020). The proximity of these networks to one another likely encumbers efforts to dissociate g from language based solely on lesion-deficit mapping because even a relatively small lesion can affect both of these networks. Such efforts are further complicated by the high degree of inter-individual variability in the anatomical locations of these functional networks, which has been demonstrated by both electrical stimulation mapping and functional connectivity fMRI studies (Ojemann 1983; Gordon et al. 2017; Marek and Dosenbach 2018). Thus, future work is still necessary discern causal neural systems for g and language on the basis of lesion-deficit mapping.

In addition to limitations noted above, it is important to note that this study included a sample with mixed lesion etiology. Although some etiologies have comparable neuropsychological outcomes (Harris et al. 2024), it would be worthwhile for larger studies to model the effects of this complex source of variance. Other important sources of unexplained variance in cognitive test scores includes individual differences in brain recovery and reorganization (i.e., neural plasticity), and health comorbidities that commonly co-occur with lesions (Camerino et al. 2020; Crockett et al. 2021; Wawrzyniak et al. 2022; Seghier and Price 2023) It will be important for future studies to compare whether maps that do and do not control for g are more accurate with respect to identifying unique anatomy for specific cognitive abilities. This could be done, for example, by predicting domain-specific cognitive deficits (i.e., from a bifactor model) in a new cohort using the lesion-deficit maps provided by this study. Lastly, it is important to recognize that the key confounds to account for in any lesion-deficit mapping study depend upon the goal of that study. The present study suggests that g is often worthy of consideration as a key confound if the goal of that study is to dissociate anatomy for different higher-order cognitive functions.

In conclusion, we present evidence for the influence of g in lesion-deficit mapping of multiple domain-specific cognitive abilities. These findings carry implications for future lesion-deficit mapping studies of cognition and provide a template for how the field can better delineate the “natural joints” of cognition using formal modeling of cognitive abilities.

## Acknowledgements

This work was supported by the National Institute of Mental Health (grant number 2T32MH019113-29A1, 1 P50 MH094258, 1 R21 MH120441-01), the National Institute of Neurological Disease and Stroke (grant number 1 R01 NS114405-01, NS095741), the National Institute of General Medical Sciences (grant number T32GM108540), the National Institutes of Health (grant number 1S10RR028821-01), and the University of Iowa Roy J. and Lucille A. Carver College of Medicine, the Kiwanis Foundation, the Ministry of Health, Italy (grant number RF-2008 -12366899, RF-2019-12369300); BIAL foundation grant (grant number 361/18); H2020 European School of Network Neuroscience (euSNN); H2020 Visionary Nature Based Actions For Heath, Wellbeing & Resilience in Cities (VARCITIES); European Union (ERC-2022-SYG NEMESIS grant number 101071900).

## Abbreviations

LBM: Lesion-Behavior Mapping
sLNM: Structural Lesion Network Mapping
fLNM: Functional Lesion Network Mapping
WAIS: Wechsler Adult Intelligence Scale
IBBM: Iowa Brain-Behavior Modeling
PLSR: Partial Least Squares Regression

