## Supplementary Material for "Cognition Disconnected: The Influence of Domain-General Cognition (g) in Lesion-Deficit Mapping"

### PLSR model visualization methods

For each PLSR model, the sign of the cognitive factor (i.e., the behavioral input into the PLSR model) was flipped so that positive model weights were associated with lower factor scores. Visualizations in figures and downstream analyses were based on the positive normalized regression weights to allow for a lesion-deficit interpretation of the results. We used a False Discovery Rate (FDR) correction (corrected alpha = 0.05) to threshold the model weights for visualization whenever possible. If no model weights survived the FDR correction, we used an alternative threshold to facilitate visual comparison of the correlated factors- and bifactor-based PLSR models: if one map in a given cognitive domain and predictor modality (e.g., correlated factors version of LBM of verbal learning/memory) had model weights that did not survive FDR correction, but its counterpart from the other psychometric model (e.g., the bifactor version of LBM of Gl) did, we applied the model weight threshold from the latter to the former so that the relative importance of the model features could be more easily compared across the maps. If neither map in a given cognitive domain and predictor modality had model weights that survived FDR correction, we applied a threshold of 0.2 to both sets of normalized voxel weights.

### Additional information on structural equation modeling procedures

We built the baseline model (Model 1) based upon previously published work in a smaller subsample of the Iowa Registry[1] in which observed variables were assigned to load on to g and the following factors based on the CHC taxonomy: crystalized intelligence (Gc), visuospatial ability (Gv), learning/memory (Gl), and processing speed (Gs). Tests that were not included in prior work were fit into the baseline model using prior work and the CHC model[2] as a guide where possible: Boston Naming Test loaded on Gc, Benton Facial Recognition Test and the Matrix Reasoning subtest from the Wechsler Adult Intelligence Scales (WAIS) loaded on Gv, and the Symbol Search subtest from the WAIS loaded on Gs. Because multiple new tests of language were included (Boston Naming Test from the Boston Diagnostic Aphasia Examination (BDAE), and Token Test, Sentence Repetition and Controlled Oral Word Association from the Multilingual

Aphasia Examination (MAE)), we also modeled a language factor (LANG) using these tests in Model 1. We then tested for the added value of modeling the covariance between WAIS subtests that belong to the same composite score domain (Model 2), the covariance among the domain-specific factors (i.e., removing orthogonality constraints on domain-specific factors; Model 3), and the following hypothesized cross-loadings that have not been previously tested (Model 4): Benton Visual Retention Test loading on Gl given the memory aspect of the task demands, COWA loading on Gs given the speed demands of the test, and WRAT loading on LANG due to its reading demands. We then examined the local fits (i.e., the statistical significance of parameter estimates) of the best-fitting model, and re-fit that model after removing parameters that did not have an associated *p*-value of less than 0.05. This final model was used to perform lesion-deficit mapping and calculate model-based reliability estimates for each latent factor using the omega-hierarchical coefficient as calculated by the compRelSEM function in R's lavaan package.

### Additional structural equation modeling results

Model 1 (the baseline model) provided a somewhat poor fit to the data (Robust CFI = 0.888, Robust TLI = 0.862, RMSEA = 0.079, SRMR = 0.085). The addition of the covariances between WAIS subtests belonging to the same composite score domain (Model 2) resulted in a non-positive definite variance-covariance matrix, and thus was rejected in favor of Model 1. The addition of non-orthogonal covariances among domain-specific factors (Model 3; Robust CFI = 0.931, Robust TLI = 0.911, Robust RMSEA = 0.063, SRMR = 0.054) fit the data significantly better than Model 1 ($X^2$ Difference = 142.99, $p < 0.001$), and thus Model 3 was retained over Model 1. The addition of hypothesized cross-loadings (Model 4; Robust CFI = 0.947, Robust TLI = 0.930, Robust RMSEA = 0.056, SRMR = 0.048) fit the data better than Model 3, and was retained over Model 3 ($X^2$ Difference = 82.72, $p < 0.001$). We re-fit Model 4 after removing the following non-significant parameters: covariances between Gc and Gs, Gc and LANG, Gv and Gl, and Gs and LANG. The final, re-fit model provided a good to excellent fit to the data (Robust CFI = 0.948, Robust TLI = 0.933, Robust RMSEA = 0.055, SRMR = 0.054) and did not statistically significantly differ in model fit to the data compared to the original Model 4 ($X^2$ Difference = 4.97, $p < 0.547$).

### Methods for LBM, sLNM, and fLNM of individual tests

To test whether g's influence on LBM and LNM was limited to the level of factor scores, we created mass-univariate lesion-deficit maps in which individual tests were the outcome variables rather than the cognitive factor scores in the primary analyses. We compared mass-univariate models with and without regressing out g. We did so for one test from each of the non-g domains, using the test with the highest factor loading in each domain in the Iowa correlated factors model: Wechsler Adult Intelligence Scale (WAIS) Block Design for visuospatial ability, Rey Auditory-Verbal Learning Test (RAVLT) Trial 5 for verbal learning/memory, WAIS Symbol Search for processing speed, and Token Test for language. Because each test was used to model the original g scores in the primary structural equation modeling results, we recalculated the g factor for the purposes of mapping each of the tests described above. Specifically, we re-estimated g after removing the g loading of the test being mapped. For example, to perform lesion-deficit mapping of Block Design while controlling for g, we first calculated g after removing the loading of Block Design onto g in the bifactor model. This g estimate was thus created without involving the Block Design test data, thus keeping the confound variable separate from the dependent variable; this was the g estimate used as a confound for the LBM, sLNM, and fLNM analyses in which we mapped Block Design while controlling for g. Each model was fit using data from participants who completed the test in question (i.e., we omitted missing data), resulting in different sample sizes for the Block Design ($n = 467$), Token Test ($n = 372$), RAVLT Trial 5 ($n = 459$), and Symbol Search ($n = 281$) models.

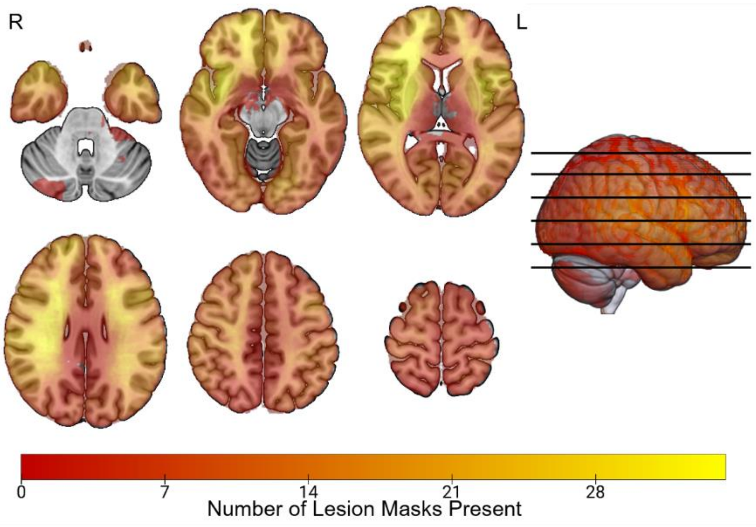


**Supplementary Figure 1 Lesion Overlap Map.** The spatial distributions of the lesion masks is depicted. Voxel values indicate the number of lesion masks present in that location.

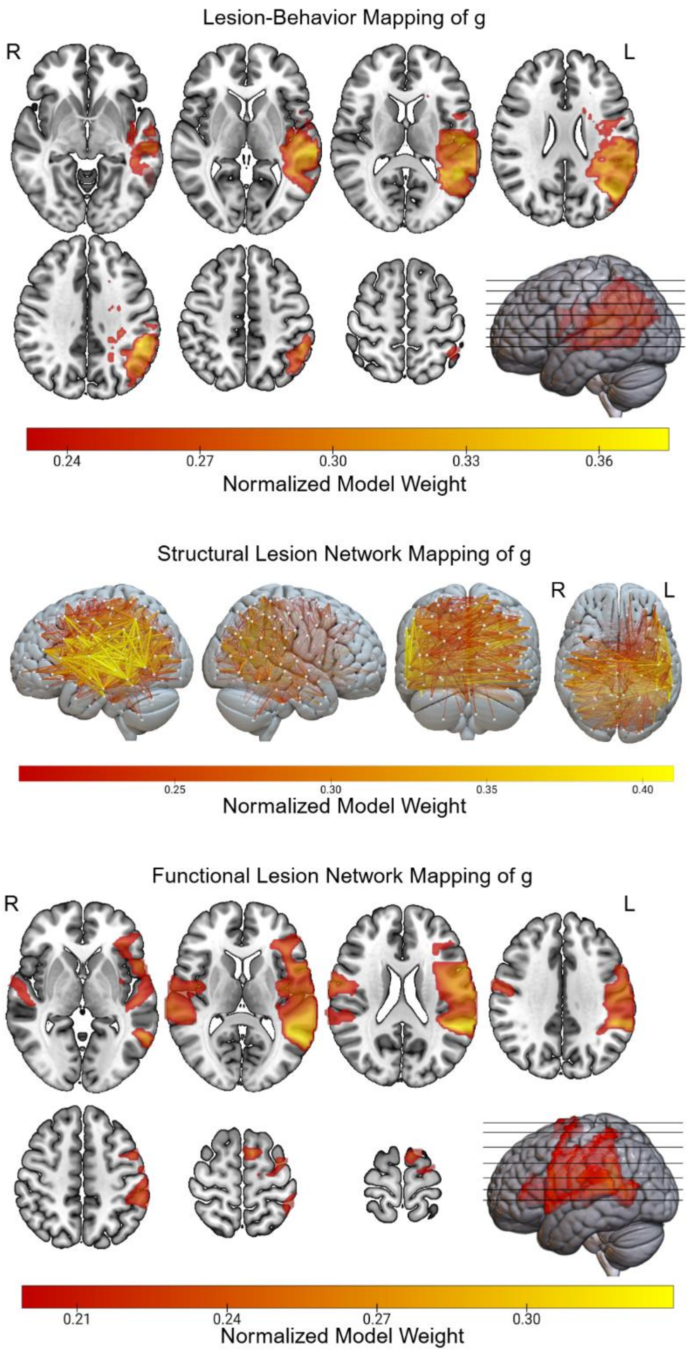


**Supplementary Figure 2 sLNM of g Controlling for Lesion Chronicity at Behavioral Testing.** The sLNM of g was performed again with lesion chronicity at behavioral testing (determined as the time of administration of the first set of WAIS subtests) as the covariate in the mass-univariate model.

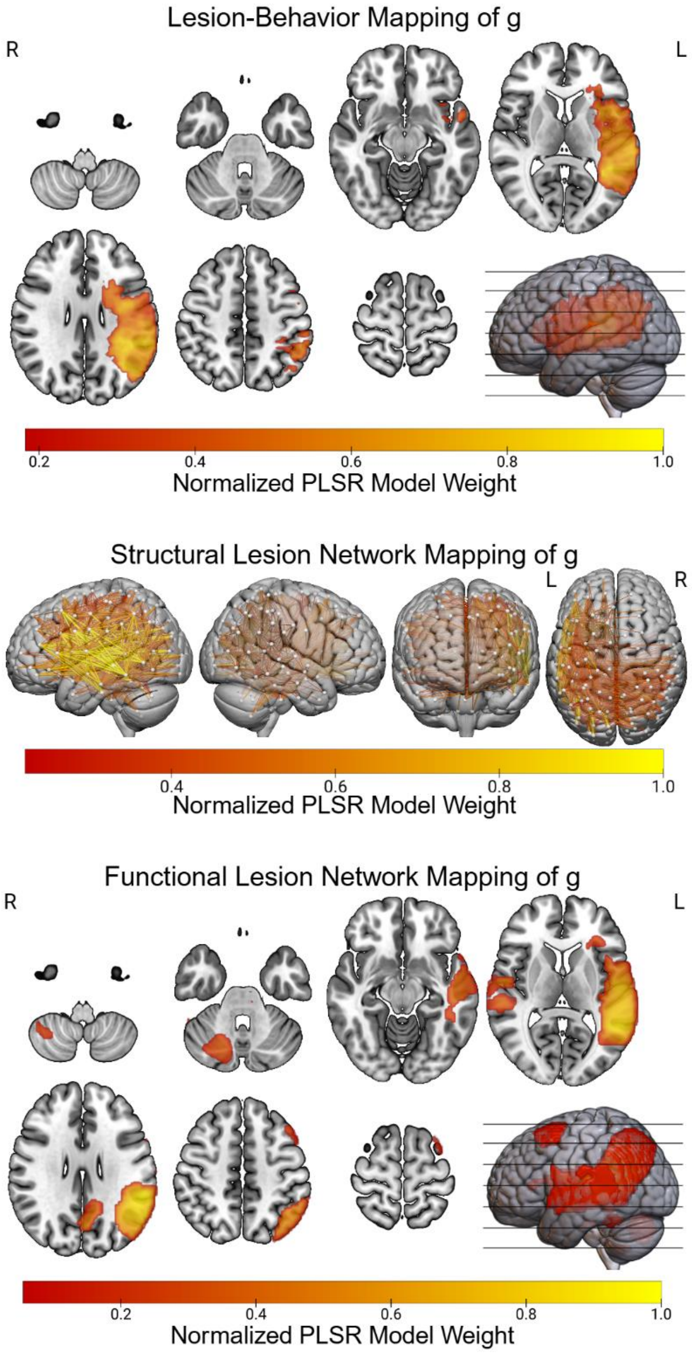


**Supplementary Figure 3 PLSR-Based Lesion-Deficit Mapping of g**. To complement the mass-univariate analyses, PLSR was used to generate LBM, sLNLM, and fLNM results for the g factor from the bifactor psychometric model shown in Figure 2.

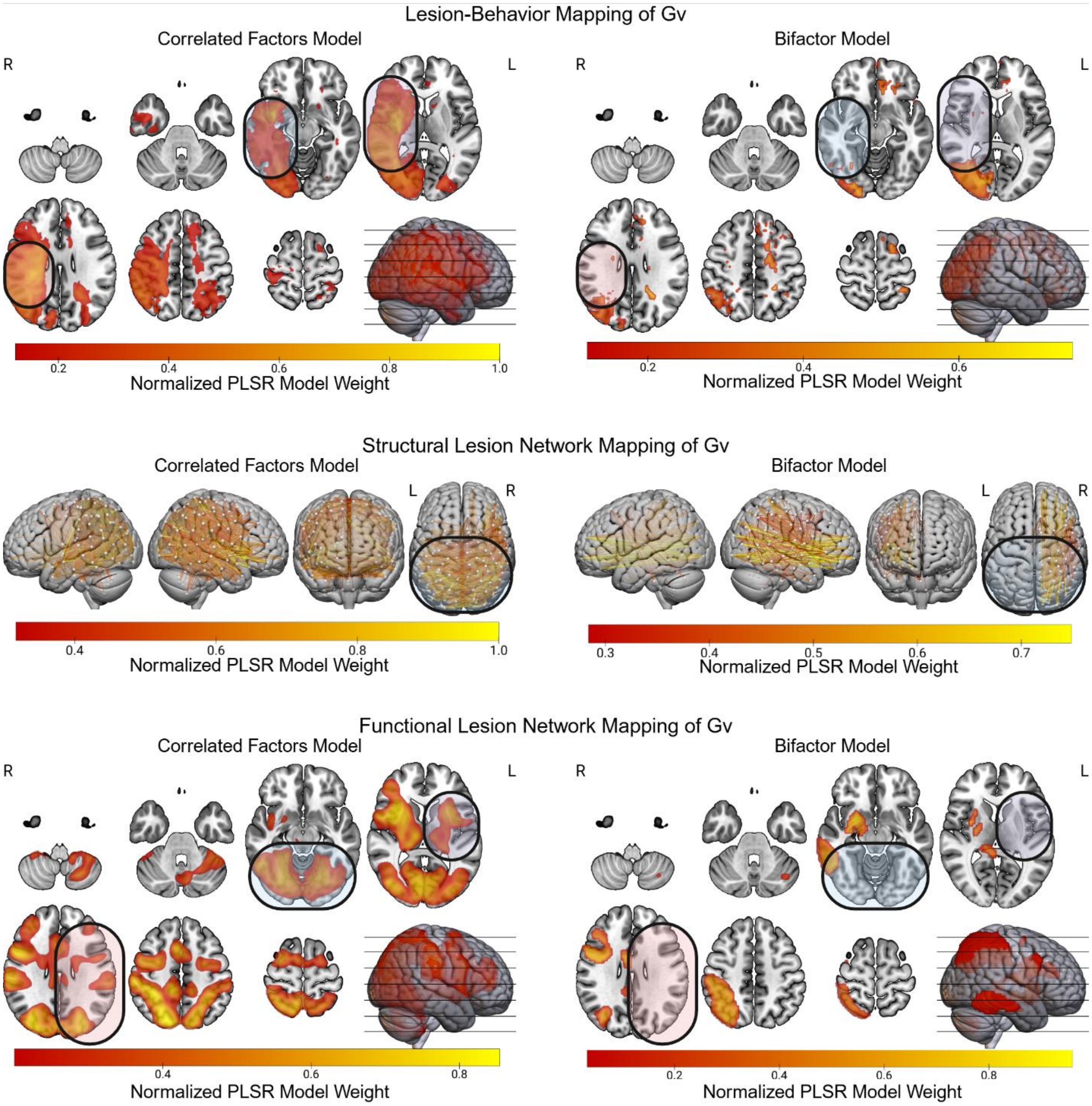


**Supplementary Figure 4 PLSR-Based Lesion-Deficit Mapping of Visuospatial Ability**. To complement the mass-univariate analyses, PLSR was used to generate LBM, sLNLM, and fLNM results for the visuospatial ability factors from the psychometric models.

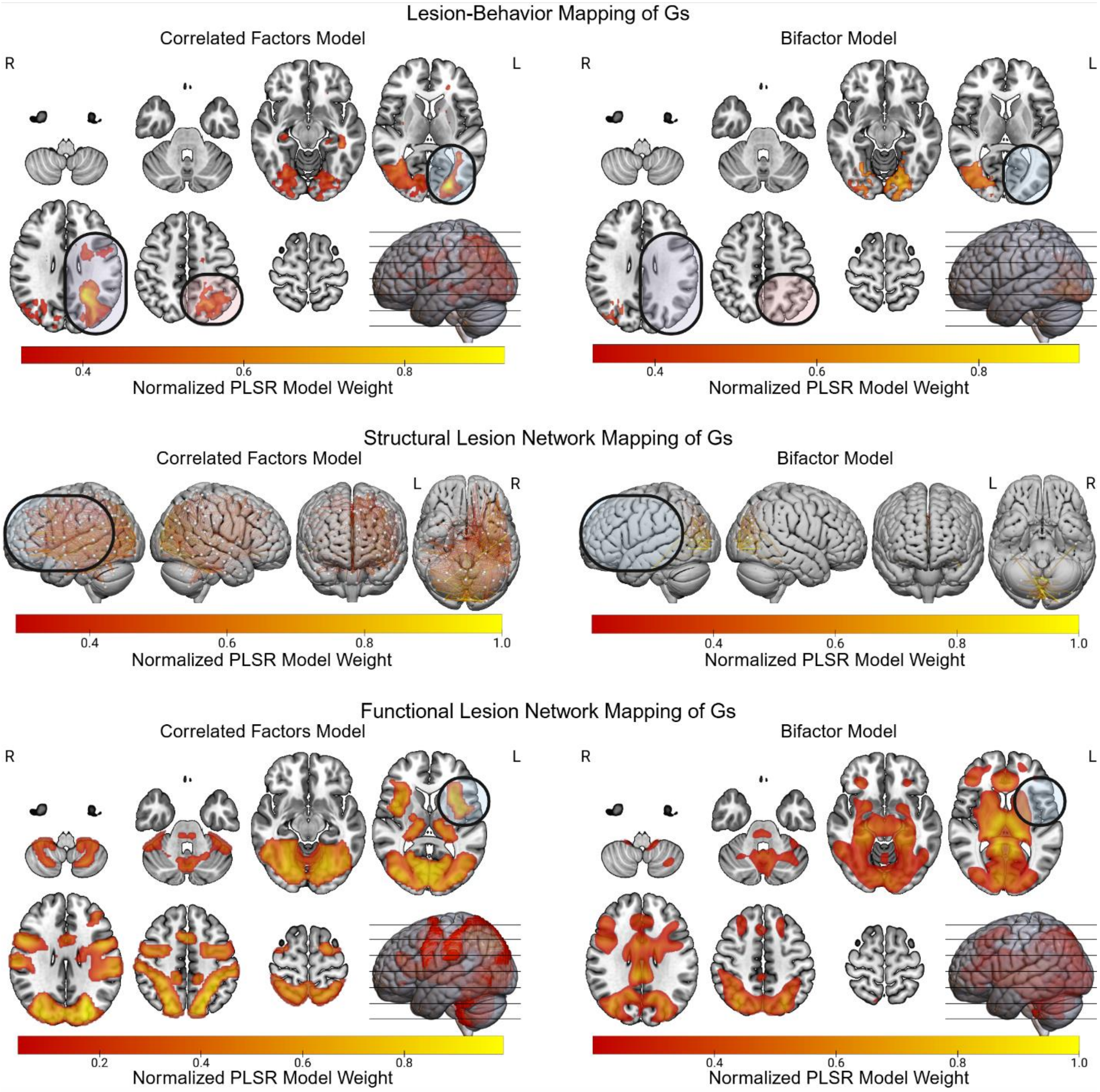


**Supplementary Figure 5 PLSR-Based Lesion-Deficit Mapping of Processing Speed**. To complement the mass-univariate analyses, PLSR was used to generate LBM, sLNLM, and fLNM results for the processing speed factors from the psychometric models.

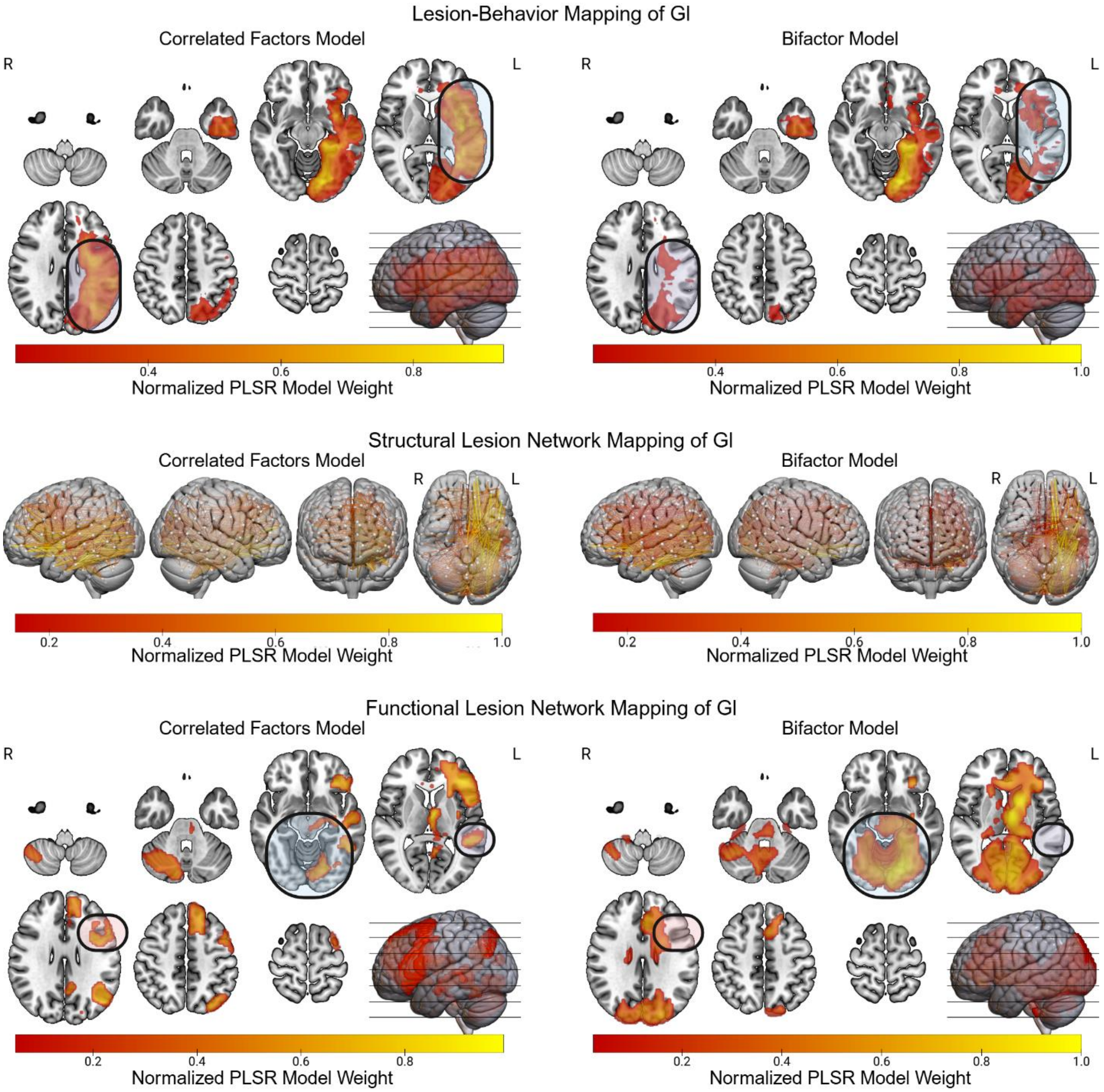


**Supplementary Figure 6 PLSR-Based Lesion-Deficit Mapping of Verbal Learning/Memory**. To complement the mass-univariate analyses, PLSR was used to generate LBM, sLNLM, and fLNM results for the verbal learning/memory factors from the psychometric models.

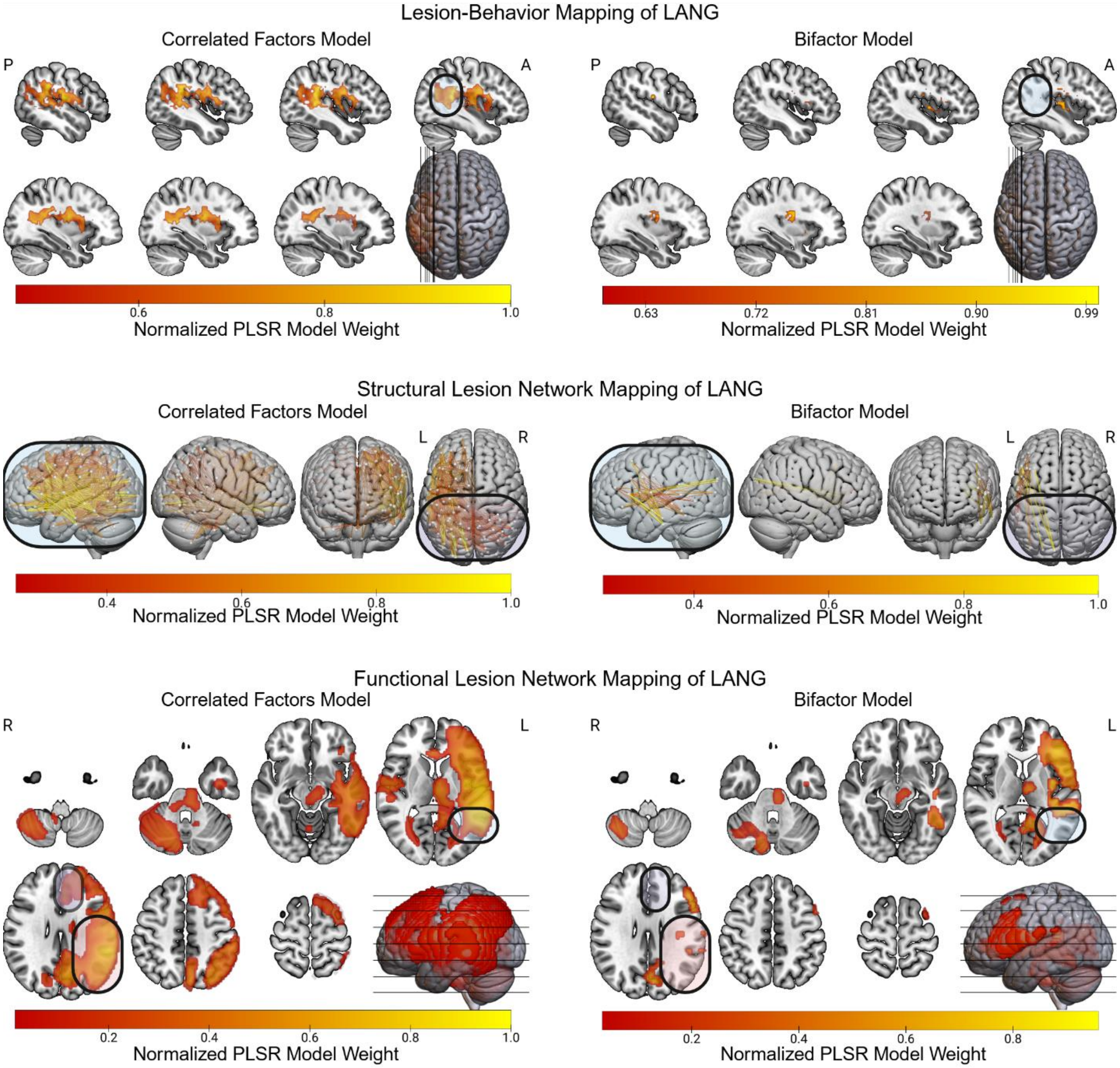


**Supplementary Figure 7 PLSR-Based Lesion-Deficit Mapping of Language**. To complement the mass-univariate analyses, PLSR was used to generate LBM, sLNLM, and fLNM results for the language factors from the psychometric models.

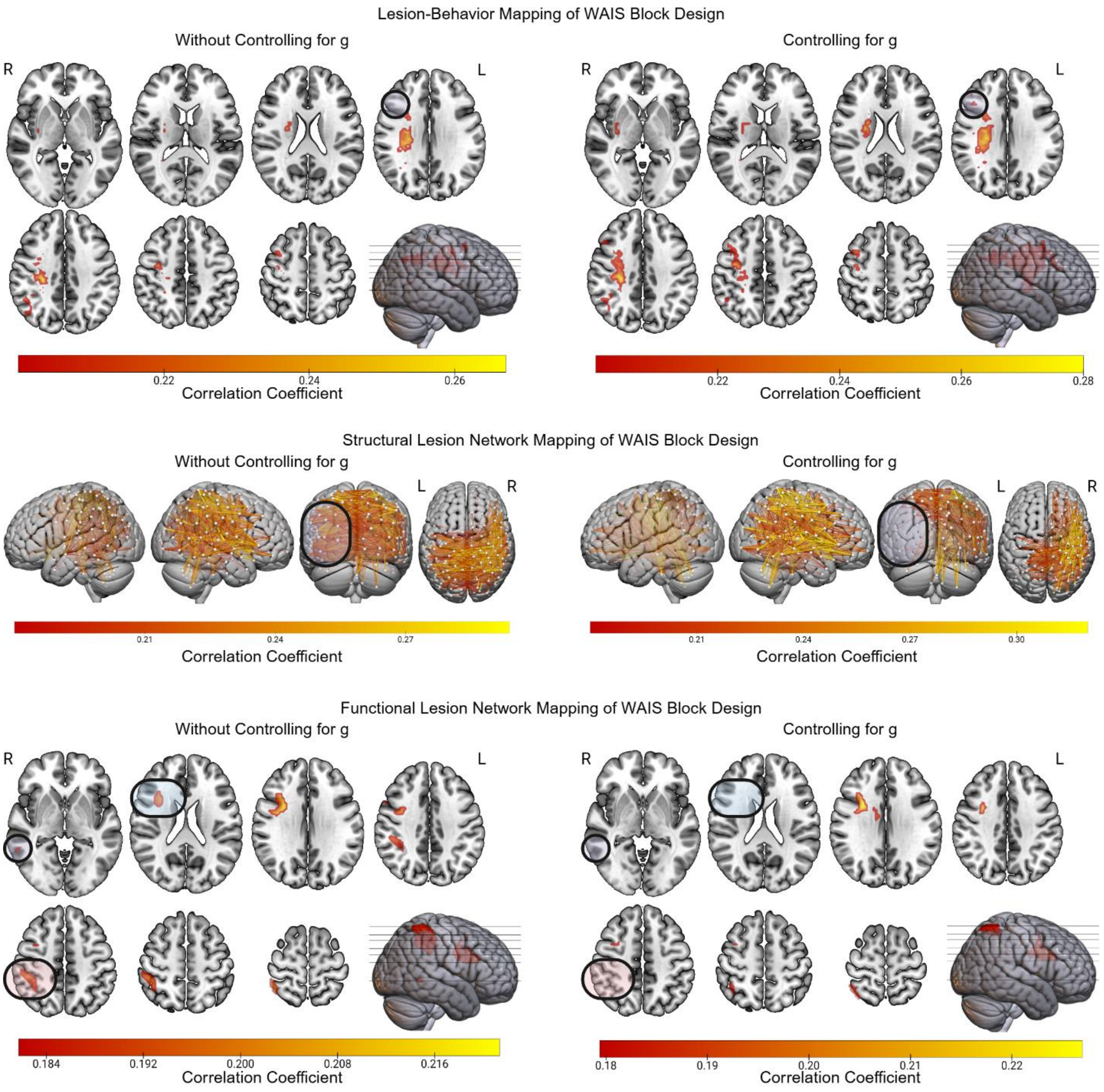


**Supplementary Figure 7 LBM and LNM of WAIS Block Design.** Lesion-Behavior Mapping, and Structural and Functional Lesion Network Mapping results are presented for the Wechsler Adult Intelligence Scale (WAIS) Block Design subtest, both with and without controlling for a version of domain-general cognition (g) derived without the use of the Block Design data (see supplemental methods). Differences between the two sets of results are highlighted.

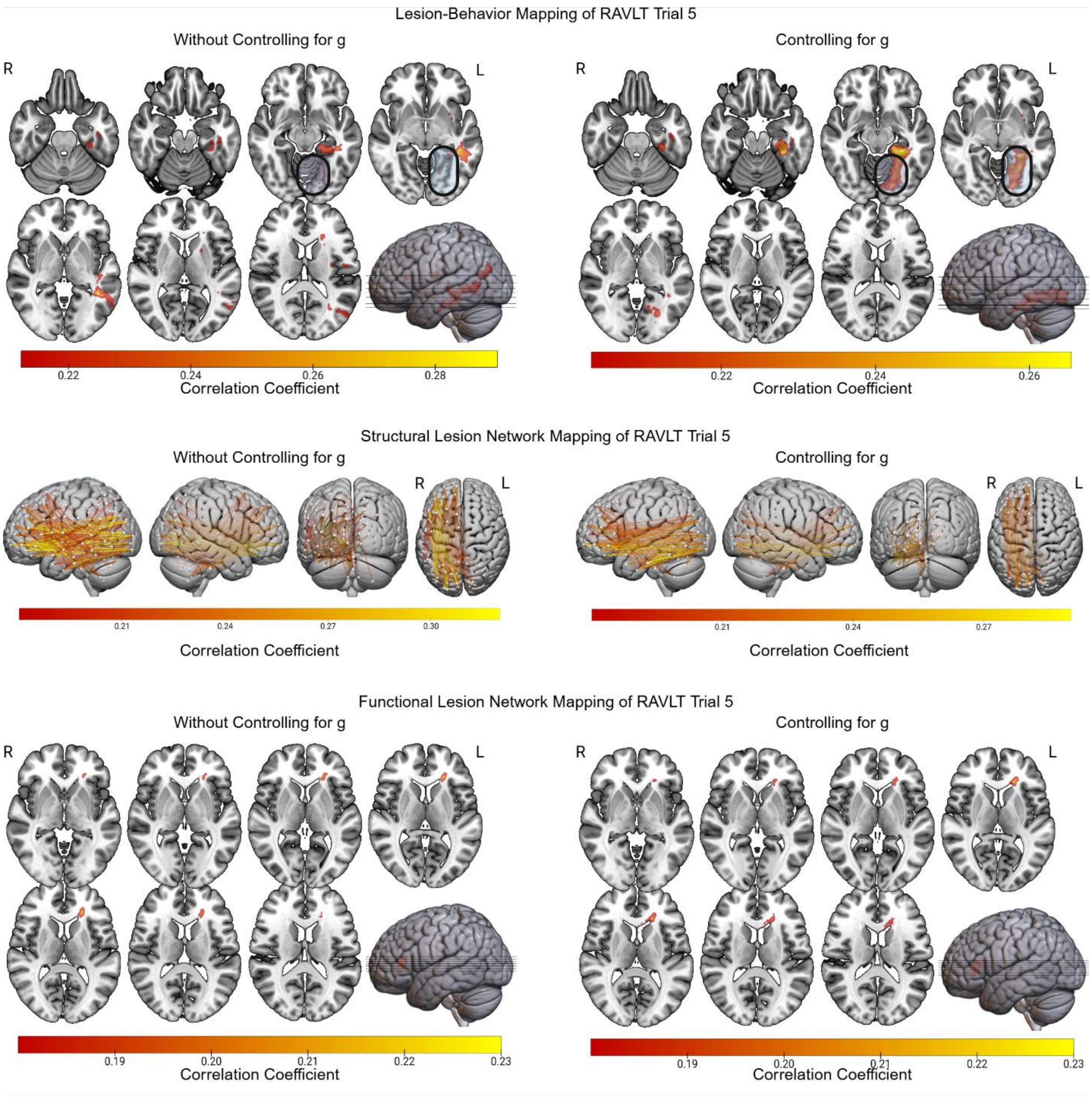


**Supplementary Figure 8 LBM and LNM of RAVLT Trial 5.** Lesion-Behavior Mapping, and Structural and Functional Lesion Network Mapping results are presented for the Rey Auditory-Verbal Learning (RAVLT) Trial 5 subtest, both with and without controlling for domain-general cognition (g), which was derived without the use of the RAVLT Trial 5 data.

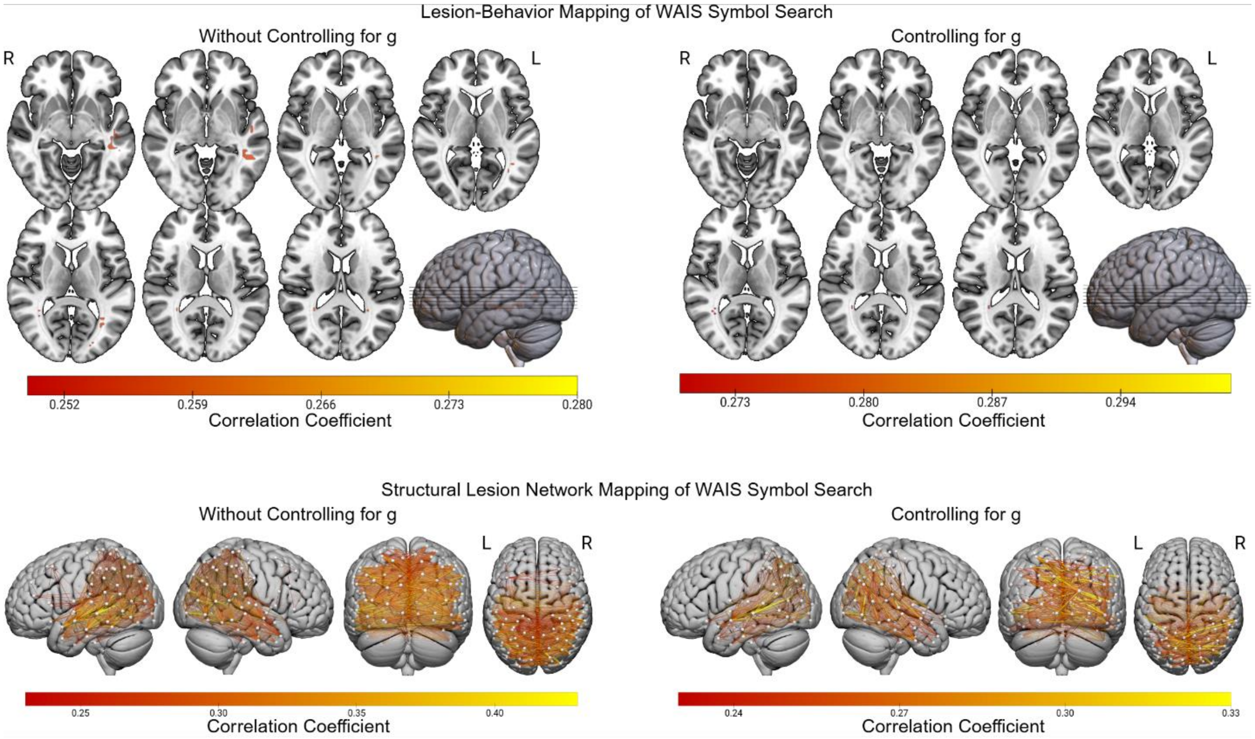


**Supplementary Figure 9 LBM and LNM of WAIS Symbol Search.** Lesion-Behavior Mapping, and Structural Lesion Network Mapping results are presented for the Wechsler Adult Intelligence Scale (WAIS) Symbol Search subtest, both with and without controlling for domain-general cognition (g), which was derived without the use of the Symbol Search data. Functional Lesion Network Mapping did not yield any statistically significant results.

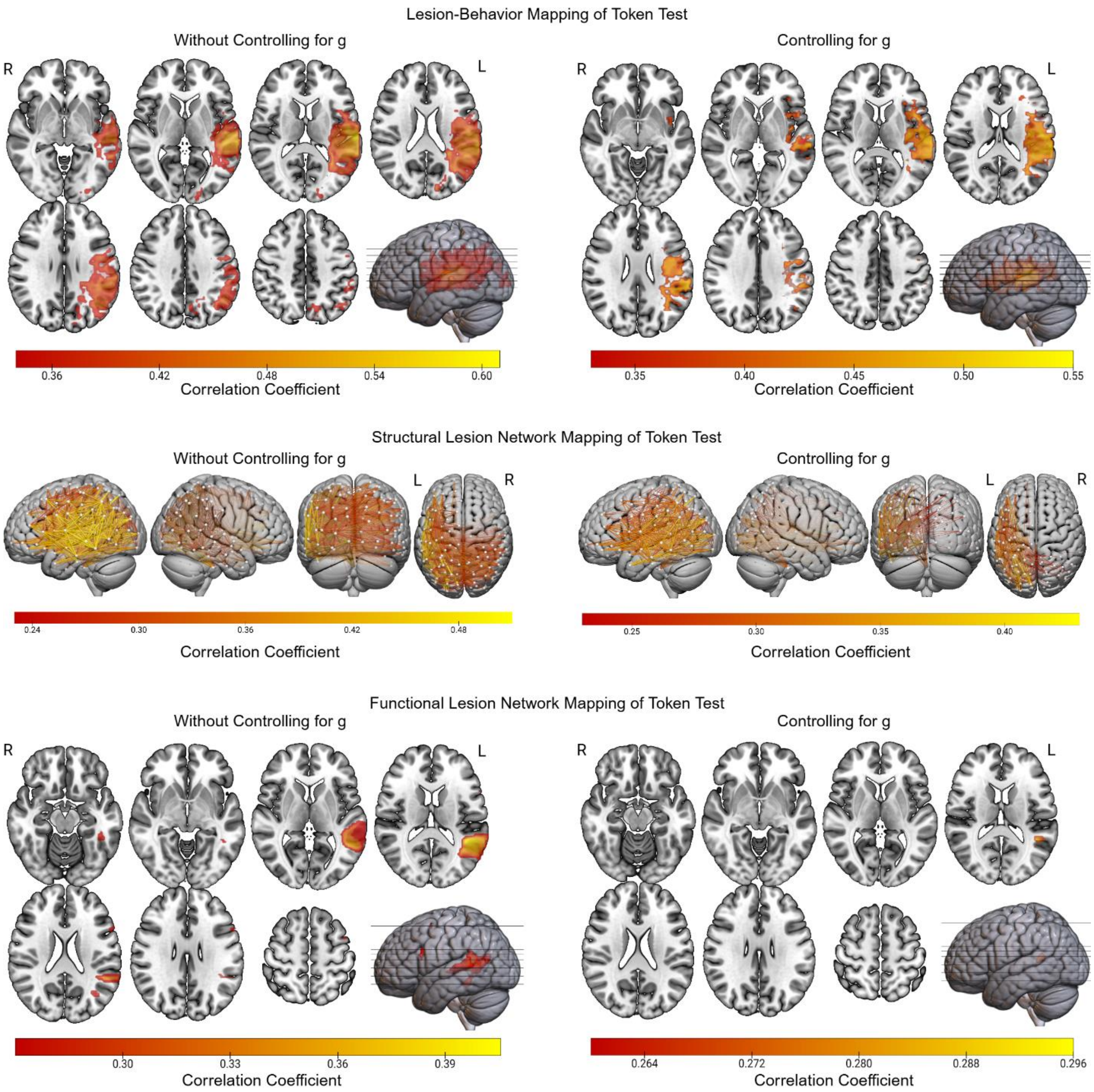


**Supplementary Figure 10 LBM and LNM of Token Test.** Structural and Functional Lesion Network Mapping results are presented for the Token Test with and without controlling for domain-general cognition (g), which was derived without the use of the Token Test data.

**Supplementary Table 1** Neuropsychological and Cognitive Tests.

| |
|---|
| WAIS Similarities* |
| WAIS Information* |
| WAIS Block Design** |
| WAIS Matrix Reasoning** |
| WAIS Digit Span* |
| WAIS Arithmetic* |
| WAIS Digit-Symbol Coding** |
| WAIS Symbol Search** |
| WRAT Word Reading* |
| Benton Judgment of Line Orientation** |
| Benton Facial Recognition Test** |
| Benton Visual Retention Test – Total Correct** |
| Rey-Osterrieth Complex Figure Test – Copy** |
| Rey-Osterrieth Complex Figure Test – Delayed Recall** |
| Rey AVLT Trial 5* |
| Rey AVLT Delayed Recall* |
| Rey AVLT Recognition Hits* |
| BDAE Boston Naming Test* |
| MAE Token Test* |
| MAE Sentence Repetition* |
| MAE Controlled Oral Word Association* |
| Trail-Making Test Part A** |
| Trail-Making Test Part B** |

Note: *Principally verbally mediated test; **Principally visually mediated test.

**Supplementary Table 3** Spatial Correlations of Mass-Univariate Lesion-Deficit Mapping Results from Each Bifactor Map and Its Correlated Factors Counterpart.

| | LBM | sLNM | fLNM | Mean (SD) Across Predictor Modalities |
|---|---|---|---|---|
| Verbal Learning/Memory | 0.918 | 0.909 | 0.822 | 0.883 (0.043) |
| Processing Speed | 0.588 | 0.692 | 0.845 | 0.708 (0.106) |
| Visuospatial Ability | 0.873 | 0.791 | 0.819 | 0.828 (0.034) |
| Language | 0.949 | 0.926 | 0.754 | 0.876 (0.087) |
| Mean (SD) Across Factors | 0.832 (0.143) | 0.830 (0.095) | 0.810 (0.034) | 0.824 (0.070) |

Spatial correlations are based on unthresholded model weights; LBM = Lesion-Behavior Mapping; sLNM = Structural Lesion Network Mapping; fLNM = Functional Lesion Network Mapping; SD = Standard Deviation.

.

**Supplementary Table 4** Anatomy Associated with Top Cluster Peaks from LBM of Cognitive Factors.

| Cluster Peak # | 1 | 2 | 3 | 4 | 5 | 6 | 7 |
|---|---|---|---|---|---|---|---|
| g | L TPOJ1 (x=-48, y=-45, z=12; r=0.383) | L A4 (x=-57, y=-27, z=9; r=0.376) | CPT O L (x=-33, y=-48, z=12; r=0.374) | L A5 (x=-66, y=-30, z=3; r=0.374) | L A4 (x=-69, y=-30, z=9; r=0.373) | DRTT L / RST L / TR S L (x=-24, y=12, z=30; r=0.233) | L 6v / SLF2 L (x=-57, y=-6, z=42; r=0.225) |
| Gl-CF | CC (x=-42, y=-48, z=0; r=0.294) | L STSvp / PAT L / AF L (x=-51, y=-45, z=-3; r=0.268) | IFOF L / ILF L (x=-42, y=-36, z=-6; r=0.267) | CC (x=-45, y=-30, z=-9; r=0.266) | L TE1p / PAT L / AF L (x=-60, y=-51, z=-3; r=0.260) | C PH L (x=-21, y=-33, z=-12; r=0.258) | L Ig / CS S L (x=-33, y=-9, z=18; r=0.246) |
| Gl-BF | L PHA2 / C PH L (x=-30, y=-36, z=-12; r=0.328) | L PHA1 (x=-24, y=-39, z=-15; r=0.327) | C PH L / C PHP L (x=-21, y=-33, z=-15; r=0.324) | L V3 (x=-21, y=-69, z=-12; r=0.304) | L V4 (x=-21, y=-75, z=-6; r=0.300) | F L (x=-33, y=-24, z=-9; r=0.212) | IFOF L (x=-30, y=9, z=-6; r=0.207) |
| Gs-CF | CC (x=-24, y=-51, z=21; r=0.276) | CC (x=-21, y=-51, z=27; r=0.268) | VOF L / IFOF L / ILF L (x=-24, y=-87, z=9; r=0.260) | CC (x=-27, y=-57, z=27; r=0.258) | MdLF L (x=-24, y=-57, z=39; r=0.257) | CC (x=-21, y=-81, z=18; r=0.249) | CC (x=-27, y=-69, z=15; r=0.243) |

| | | | | | | | |
|---|---|---|---|---|---|---|---|
| Gs-BF | R MST (x=48, y=-69, z=6; r=0.247) | CC (x=30, y=-51, z=12; r=0.240) | TR P R / IFOF R / ILF R (x=36, y=-57, z=6; r=0.237) | CC (x=27, y=-69, z=24; r=0.230) | CPT O L (x=-21, y=-81, z=12; r=0.228) | TR S R (x=21, y=-24, z=21; r=0.226) | CC (x=-18, y=-87, z=6; r=0.226) |
| Gv-CF | IFOF R / ILF R (x=36, y=-57, z=9; r=0.261) | CC (x=30, y=-60, z=9; r=0.251) | R Putamen (x=33, y=-12, z=0; r=0.250) | R PGs / SLF2 R (x=42, y=-66, z=42; r=0.250) | CC (x=30, y=-48, z=15; r=0.247) | CC (x=27, y=-45, z=21; r=0.247) | R PGs (x=45, y=-75, z=36; r=0.245) |
| Gv-BF | R Putamen (x=33, y=-12, z=0; r=0.295) | ILF R / IFOF R (x=36, y=-57, z=9; r=0.283) | EMC R (x=36, y=3, z=-9; r=0.276) | ML R (x=24, y=-27, z=24; r=0.276) | CC (x=27, y=-45, z=21; r=0.271) | ILF R (x=36, y=-63, z=9; r=0.268) | CC (x=30, y=-48, z=15; r=0.266) |
| LANG -CF | L A4 (x=-69, y=-30, z=9; r=0.522) | L PFop (x=-63, y=-24, z=15; r=0.514) | L A4 (x=-66, y=-18, z=12; r=0.488) | L STV (x=-66, y=-39, z=15; r=0.486) | L TE1a (x=-66, y=-18, z=-12; r=0.302) | CC (x=-18, y=-57, z=33; r=0.289) | L V2 (x=-24, y=-102, z=0; r=0.289) |
| LANG -BF | L A4 (x=-69, y=-27, z=9; r=0.453) | L A4 (x=-66, y=-18, z=12; r=0.43) | CC (x=-42, y=-33, z=0; r=0.403) | L STV (x=-66, y=-45, z=15; r=0.400) | L 45 (x=-39, y=27, z=3; r=0.343) | L AVI (x=-30, y=27, z=6; r=0.336) | RST L / IFOF L (x=-42, y=27, z=9; r=0.333) |

CF = Correlated Factors Mode; BF = Bifactor Model; Gv = Visuospatial Ability; Gl = Verbal Learning/Memory; Gs = Processing Speed; LANG = Language; Coordinates are in MNI152 1mm space; Anatomical descriptors were derived from the Human Connectome Project (HCP) Multimodal Parcellation Atlas Version 1 and the Harvard-Oxford Subcortical Atlas for results in cortical and sub-cortical regions, and the HCP 1065 group-averaged white matter atlas for results in white matter; w = Partial Least Squares Regression Weight Value; L = Left Hemisphere; R = Right Hemisphere.

**Supplementary Table 5** Cortical Parcel Pairs Associated with Top Model Weights from sLNM of Cognitive Factors.

| Weight # | 1 | 2 | 3 | 4 | 5 | 6 | 7 |
|---|---|---|---|---|---|---|---|
| G | L SomMot 4 and L Limbic TempPole 2 (r=0.397) | L Cont Par 1 and L Default Temp 4 (r=0.395) | L SomMot 6 and L DosAttn Post 2 (r=0.393) | L Cont Par 1 and L Cont Temp 1 (r=0.392) | L SomMot 4 and L Cont Temp 1 (r=0.391) | L SomMot 6 and L Cont Temp 1 (r=0.391) | L SomMot 4 and L Default Temp 4 (r=0.390) |
| Gl-CF | L Vis 9 and L Default PFC 5 (r=0.324) | L Vis 14 and L Default PFC 5 (r=0.322) | L Vis 2 and L Default PFC 3 (r=0.322) | L Vis 7 and LH SalVentAttn FrOperIns 2 (r=0.321) | L Vis 13 and L Default PFC 5 (r=0.320) | L Vis 14 and L Default PFC 3 (r=0.319) | L Vis 11 and L Default PFC 5 (r=0.318) |
| Gl-BF | L Vis 2 and L Limbic TempPole 2 (r=0.327) | L Vis 7 and L Limbic TempPole 2 (r=0.326) | L Vis 4 and L Limbic TempPole 2 (r=0.323) | L Default pCunPCC 1 and L Default PHC 1 (r=0.319) | L Vis 6 and L Default PHC 1 (r=0.316) | L Vis 10 and L Limbic TempPole 2 (r=0.314) | L Vis 1 and L Limbic TempPole 2 (r=0.311) |
| Gs-CF | L Vis 7 and R Vis 14 (r=0.317) | L Default Temp 4 and R Vis 12 (r=0.317) | L Default Temp 3 and R Vis 13 (r=0.313) | L Vis 7 and R Vis 12 (r=0.310) | L Default Temp 3 and R Vis 3 (r=0.309) | L Default Temp 4 and R Vis 13 (r=0.307) | L Vis 13 and R Vis 14 (r=0.304) |

| | | | | | | | |
|---|---|---|---|---|---|---|---|
| Gs-BF | L Vis 7 and R Vis 12 (r=0.328) | L Vis 7 and R Vis 14 (r=0.326) | L Vis 7 and R Vis 4 (r=0.319) | L Vis 7 and R Vis 9 (r=0.319) | L Vis 13 and R Vis 6 (r=0.317) | L Vis 13 and R Vis 12 (r=0.315) | L Vis 14 and R Vis 9 (r=0.311) |
| Gv-CF | L DorsAttn Post 8 and R Vis 5 (r=0.287) | L DorsAttn Post 9 and R Default Temp 5 (r=0.280) | L DorsAttn Post 8 and R Vis 15 (r=0.276) | L DorsAttn Post 7 and R DorsAttn Post 1 (r=0.276) | L DorsAttn Post 3 and R Vis 5 (r=0.274) | L DorsAttn Post 7 and R Vis 154 (r=0.271) | L DorsAttn Post 3 and R DorsAttn Post 2 (r=0.271) |
| Gv-BF | R Vis 12 and R Limbic OFC 2 (r=0.334) | R Vis 12 and R Cont PFCv 1 (r=0.330) | R Vis 14 and R Cont PFCv 1 (r=0.328) | R Vis 12 and R Default PFCv 1 (r=0.325) | R Vis 12 and R SalVentAttn FrOperIns 4 (r=0.322) | R Vis 6 and R Default PFCv 1 (r=0.321) | R Vis 15 and R Default PFCv 1 (r=0.320) |
| LANG-CF | L SomMot 4 and L Limbic TempPole 2 (r=0.513) | L Vis 13 and L SalVentAttn FrOperIns 2 (r=0.490) | L SomMot 2 and L SomMot 4 (r=0.487) | L Vis 13 and L Default PFC 5 (r=0.486) | L Vis 14 and L Default PFC 5 (r=0.486) | L SomMot 4 and L Default Temp 4 (r=0.482) | L Vis 11 and L Default PFC 5 (r=0.482) |
| LANG-BF | L SalVent Attn ParOper 2 and left thalamus | L Default Temp 5 and left thalamus proper (r=0.382) | L SalVentAttn ParOper 1 and left thalamus | L SomMot 5 and left thalamus proper (r=0.372) | L SomMot 4 and L Limbic TempPole | L Vis 9 and L Default PFC 5 (r=0.367) | L Vis 7 and L Default PFC 3 (r=0.365) |

| | | |
|---|---|---|
| proper (r=0.385) | proper (r=0.377) | 2 (r=0.368) |

Anatomical descriptors were derived from existing atlases spanning cortical, subcortical, and brainstem regions as described in the methods section of the primary text; CF = Correlated Factors Model; BF = Bifactor Model; Gv = Visuospatial Ability; Gl = Verbal Learning/Memory; Gs = Processing Speed; LANG = Language; r = Pearson Correlation Coefficient Value; L = Left Hemisphere; R = Right Hemisphere.

**Supplementary Table 6** Anatomy Associated with Top Cluster Peaks from fLNM of Cognitive Factors.

| Cluster Peak # | 1 | 2 | 3 | 4 | 5 | 6 | 7 |
|---|---|---|---|---|---|---|---|
| g | TR S L / SLF3 L / AF L (x=-42, y=-42, z=30; r=0.334) | CC (x=-54, y=-54, z=6; r=0.322) | L TPOJ2 (x=-45, y=-54, z=12; r=0.316) | AF L / FAT L (x=-42, y=12, z=15; r=0.271) | RST L / IFOF L (x=-42, y=27, z=9; r=0.270) | CB R (x=18, y=-63, z=-27; r=0.249) | V (x=21, y=-72, z=-45; r=0.248) |
| Gl-CF | L PuM / OR L (x=-12, y=-33, z=0; r=0.235) | CB R (x=21, y=-75, z=-21; r=0.224) | CC (x=-9, y=21, z=15; r=0.218) | CB R (x=39, y=-60, z=-24; r=0.218) | IFOF L (x=-42, y=-42, z=-9; r=0.209) | CS A L (x=-24, y=33, z=6; r=0.207) | V (x=15, y=-75, z=-42; r=0.200) |
| Gl-BF | L PuM / OR L (x=-12, y=-33, z=0; r=0.284) | CC (x=-9, y=24, z=12; r=0.236) | CB R (x=27, y=-78, z=-21; r=0.231) | CB R (x=9, y=-81, z=-18; r=0.226) | L V1 (x=31, y=22, z=26; r=0.221) | CB R (x=45, y=-60, z=-24; r=0.217) | L PHA3 (x=-30, y=-39, z=-18; r=0.213) |
| Gs-CF | F L (x=-21, y=-36, z=6; r=0.259) | L PuM (x=-15, y=-30, z=3; r=0.254) | CC (x=-18, y=-66, z=36; r=0.246) | OR L (x=-30, y=-30, z=0; r=0.245) | L IP1 (x=-27, y=-69, z=36; r=0.241) | L VVC (x=-33, y=-33, z=-30; r=0.241) | F R (x=24, y=-36, z=6; r=0.240) |
| Gs-BF | F L (x=-21, y=-36, z=6; r=0.370) | R PuM (x=21, y=-33, | CC (x=0, y=9, z=18; r=0.287) | CC (x=-18, y=-12, z=30; r=0.217) | CPT P R (x=9, y=-30, z=-27; r=0.195) | - | - |

| | | | | | | | |
|---|---|---|---|---|---|---|---|
| | | z=6; r=0.362) | | | | | |
| Gv-CF | C PHP R (x=18, y=-60, z=39; r=0.193) | R TE2a (x=57, y=-42, z=-27; r=0.192) | CC (x=3, y=6, z=21; r=0.186) | CC (x=-18, y=-12, z=30; r=0.182) | CPT F R / CS S R / TR S R / FAT R (x=27, y=6, z=24; r=0.182) | CC (x=9, y=-6, z=27; r=0.181) | - |
| Gv-BF | R PuM / OR R (x=18, y=-33, z=6; r=0.276) | V (x=-9, y=-75, z=-39; r=0.246) | CC (x=6, y=-6, z=24; r=0.245) | C PHP R (x=21, y=-60, z=33; r=0.242) | C PHP R (x=18, y=-63, z=39; r=0.240) | CC (x=9, y=-75, z=45; r=0.234) | ICP L (x=-9, y=-66, z=-30; r=0.231) |
| LANG-CF | AF L (x=-42, y=-45, z=9; r=0.354) | IFOF L (x=-45, y=30, z=6; r=0.314) | IFOF L (x=-33, y=36, z=-3; r=0.303) | CB R (x=18, y=-69, z=-24; r=0.298) | MCP (x=-9, y=-18, z=-33; r=0.272) | L TE2p (x=-48, y=-51, z=-24; r=0.243) | L TE2p (x=-51, y=-39, z=-24; r=0.236) |
| LANG-BF | CC (x=-42, y=-42, z=6; r=0.254) | PAT L / AF L (x=-42, y=-48, z=15; 2=0.237) | CB R (x=18, y=-69, z=-24; r=0.219) | L TE1p (x=-48, y=-57, z=-21; r=0.214) | - | - | - |

CF = Correlated Factors Mode; BF = Bifactor Model; Gv = Visuospatial Ability; Gl = Verbal Learning/Memory; Gs = Processing Speed; LANG = Language; Coordinates are in MNI152 1mm space; Anatomical descriptors were derived from the Human Connectome Project (HCP) Multimodal Parcellation Atlas Version 1 and the Harvard-Oxford Subcortical Atlas for results in cortical and sub-cortical regions, and the HCP 1065 group-averaged white matter atlas for results in white matter; r = Pearson Correlation Coefficient Value; L = Left Hemisphere; R = Right Hemisphere.

**Supplementary Table 7** PLSR-Based Modeling Results.

| Outcome Variable | Predictor Modality | Inferential Model Proportion Variance Explained |
|---|---|---|
| g (residualized on log lesion volume) | LBM | $0.198^{***}$ |
| | sLNM | $0.215^{***}$ |
| | fLNM | $0.191^{***}$ |
| Verbal Learning/Memory – Correlated Factors (residualized on log lesion volume) | LBM | $0.130^{*}$ |
| | sLNM | $0.204^{***}$ |
| | fLNM | $0.124^{***}$ |
| Processing Speed – Correlated Factors (residualized on log lesion volume) | LBM | 0.199 |
| | sLNM | $0.121^{**}$ |
| | fLNM | $0.043^{***}$ |
| Visuospatial Ability – Correlated Factors (residualized on log lesion volume) | LBM | 0.094 |
| | sLNM | $0.075^{**}$ |
| | fLNM | $0.227^{*}$ |
| Language – Correlated Factors (residualized on log lesion volume) | LBM | $0.413^{***}$ |
| | sLNM | $0.361^{***}$ |
| | fLNM | $0.269^{***}$ |
| Verbal Learning/Memory – Bifactor (residualized on log lesion volume) | LBM | 0.106 |
| | sLNM | $0.126^{**}$ |
| | fLNM | $0.078^{***}$ |

| | | |
|---|---|---|
| Processing Speed – Bifactor (residualized on log lesion volume) | LBM | 0.236* |
| | sLNM | 0.177** |
| | fLNM | 0.114*** |
| Visuospatial Ability– Bifactor (residualized on log lesion volume) | LBM | 0.396*** |
| | sLNM | 0.182*** |
| | fLNM | 0.167*** |
| Language – Bifactor (residualized on log lesion volume) | LBM | 0.257*** |
| | sLNM | 0.245*** |
| | fLNM | 0.184*** |
